# KLF4 binding is involved in the organization and regulation of 3D enhancer networks during acquisition and maintenance of pluripotency

**DOI:** 10.1101/382473

**Authors:** Dafne Campigli Di Giammartino, Andreas Kloetgen, Alexander Polyzos, Yiyuan Liu, Daleum Kim, Dylan Murphy, Abderhman Abuhashem, Paola Cavaliere, Boaz Aronson, Veevek Shah, Noah Dephoure, Matthias Stadtfeld, Aristotelis Tsirigos, Effie Apostolou

**Author notes:** These authors contributed equally (Dafne Campigli Di Giammartino, Andreas Kloetgen, Alexander Polyzos).

## Abstract

Cell fate transitions are accompanied by global transcriptional, epigenetic and topological changes driven by transcription factors (TFs), as is strikingly exemplified by reprogramming somatic cells to pluripotent stem cells (PSCs) via expression of OCT4, KLF4, SOX2 and cMYC. How TFs orchestrate the complex molecular changes around their target gene loci in a temporal manner remains incompletely understood. Here, using KLF4 as a paradigm, we provide the first TF-centric view of chromatin reorganization and its association to 3D enhancer rewiring and transcriptional changes of linked genes during reprogramming of mouse embryonic fibroblasts (MEFs) to PSCs. Inducible depletion of KLF factors in PSCs caused a genome-wide decrease in the connectivity of enhancers, while disruption of individual KLF4 binding sites from PSC-specific enhancers was sufficient to impair enhancer-promoter contacts and reduce expression of associated genes. Our study provides an integrative view of the complex activities of a lineage-specifying TF during a controlled cell fate transition and offers novel insights into the order and nature of molecular events that follow TF binding.

## INTRODUCTION

The identity of each cell type is determined by a unique gene expression program as well as a characteristic epigenetic landscape and three-dimensional (3D) chromatin topology. All of these features are under the control and constant supervision of a network of critical transcription factors (TFs), known as master regulators of cell identity^1, 2^. Although the ability of master regulators to maintain or change cell identity is well accepted, the underlying mechanisms remain poorly understood.

Somatic cell reprogramming into induced pluripotent stem cells (iPSCs) by the so-called Yamanaka factors OCT4, KLF4, SOX2 and cMYC (OKSM) offers a tractable system to study the molecular mechanisms of cell fate determination and the roles and activities of each reprogramming TF^3, 4^. Research over the last decade started dissecting on a genome-wide level the transcriptional and epigenetic changes that result in successful erasure of somatic identity and establishment of pluripotency^5–7^. Distinct or synergistic roles of the reprogramming TFs as well as specific direct and indirect mechanisms for coordinating these molecular changes have been proposed ^8–14^. In addition to the transcriptional and epigenetic changes, recent studies utilizing targeted or global chromatin conformation capture techniques revealed that the 3D chromatin topology differs between somatic and pluripotent stem cells (PSCs) and is largely reset during reprogramming^15–21^. However, the principles of chromatin reorganization during iPSC generation, its association with enhancer and gene activity and the involvement of TFs in these processes have only started to be explored.

Current models regarding the role of reprogramming TFs in 3D chromatin organization are mostly based on computational analyses of 4C or HiC datasets, which reveal a strong enrichment of OKS binding around long-range interactions in PSCs and during reprogramming ^15–17, 21^. For KLF4, an architectural function is also supported by experimental evidence. In fact, KLF4 depletion abrogates loops at specific genomic loci such as the *Pou5f1* (*Oct4*) locus in the context of mouse PSCs^18^, and the *HOPX* gene in human epidermal keratinocytes^22^. In addition, depletion of the related factor KLF1 disrupts selected long-range interactions in the context of erythropoiesis^23, 24^. These findings establish a link between TF binding and chromatin architecture and suggest that OKS, and particularly KLF4, may actively orchestrate long-range chromatin interactions during reprogramming in order to establish and maintain the pluripotent transcriptional program. To directly test this possibility in a genome-wide manner, we captured the dynamic KLF4-centric topological reorganization during the course of reprogramming and determined the relationships with epigenetic and transcriptional changes. To do so, we used a well-characterized system to reprogram mouse embryonic fibroblasts (MEFs) to iPSCs ^14, 25^ and applied genome-wide assays that map KLF4 binding (ChIP-seq), chromatin accessibility (ATAC-seq), enhancer and gene activity (H3K27ac ChIP-seq and RNA-seq), enhancer connectivity (H3K27ac HiChIP) as well as KLF4-centric chromatin looping (KLF4 HiChIP) at different stages during acquisition of pluripotency (Fig.1a, top panel). Integrative analysis of our results generated a reference map of stage-specific chromatin changes around KLF4 bound loci and established strong links with enhancer rewiring and concordant transcriptional changes. Inducible depletion of KLF factors in PSCs or genetic disruption of KLF4 binding sites within specific PSC enhancers further supported the ability of KLF4 to function both as a transcriptional regulator and a chromatin organizer.

**Figure 1.**
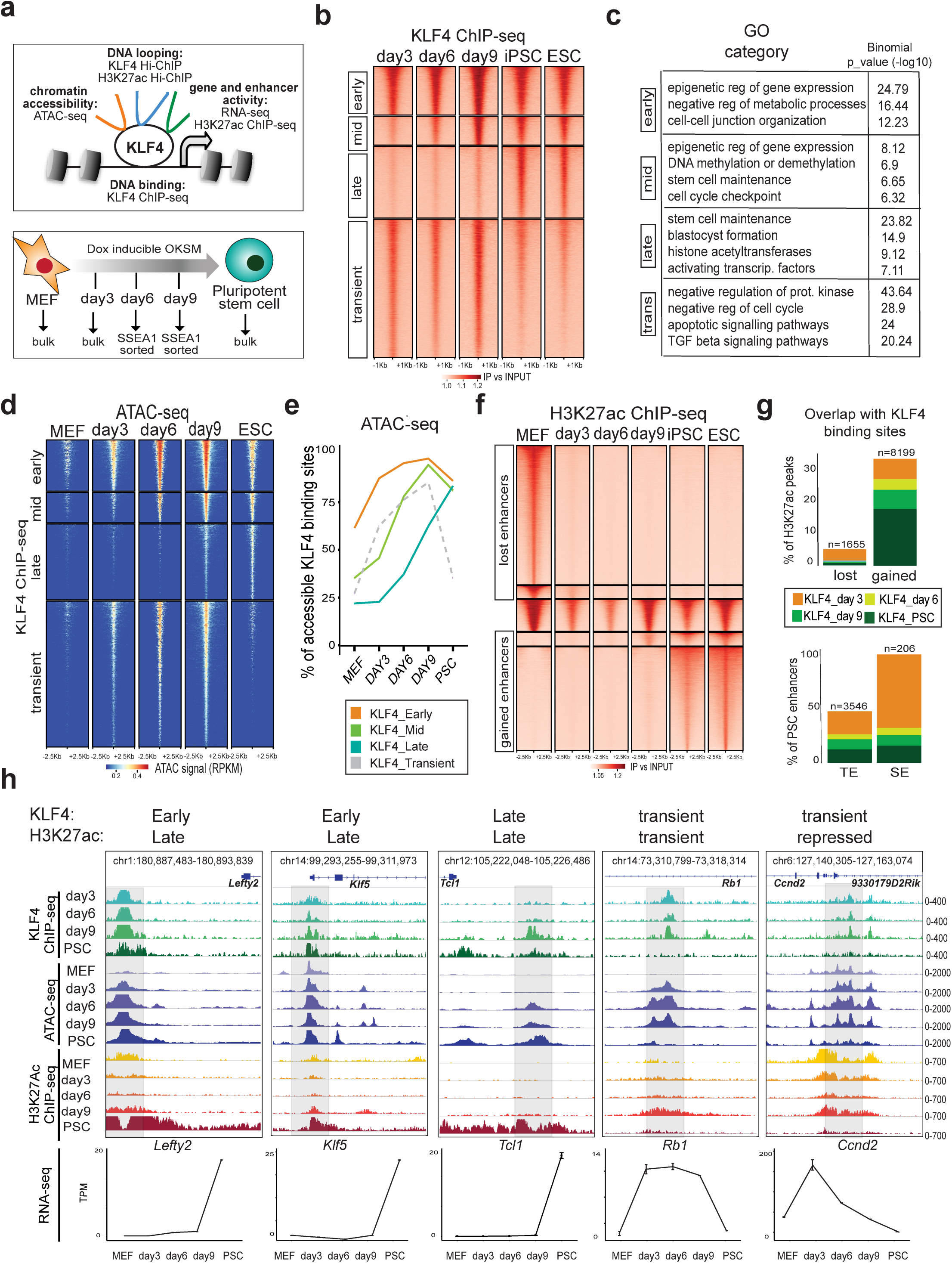
Dynamic KLF4 binding during reprogramming and association with chromatin accessibility and enhancer activity. **a** Schematic illustration of the experimental system and strategy. **b** Tornado plots of KLF4 ChIP-seq signals at different reprogramming stages clustered in four different categories: Early, Mid, Late and Transient KLF4 binding. ChIP-seq signals (fold enrichment over input) are showing 1kb upstream/downstream of peak centers. **c**, GREAT gene ontology analysis of Early, Mid, Late and Transient KLF4 target sites. **d**, Tornado plot of ATAC-seq signal ar different reprogramming stages around KLF4 binding sites (Early, Mid, Late, Transient). ATAC-seq signals are showing 2.5kb upstream/downstream of peak centers. RPKM (Read Per Kilobase Million). **e**, Line plots showing the percentages of KLF4 Early, Mid, Late and Transient targets that overlapped with ATAC-seq peaks (accessible regions) at each reprogramming stage **f**, Tornado plot of H3K27ac ChIP-seq signal showing MEF peaks, PSC peaks and constant peaks at each reprogramming stage. ChIP-seq signals (fold enrichment over input) are showing 2.5kb upstream/downstream of peak centers **g**, Bar plots showing overlap of KLF4 binding with either lost (MEF) or gained (PSC) H3K27ac peaks (top) or with typical PSC enhancers (TE) versus superenhancers (SE) (as characterized by Whyte et al., 2013) (bottom). **h**, Examples of genomic regions (see genomic coordinates) that show different kinetics of KLF4 binding and H3K27ac occupancy during reprogramming. IGV tracks for KLF4 ChIP-seq, ATAC-seq, H3K27ac ChIP-seq at each reprogramming stage are shown and the signal values are indicated on the right. The transcriptional changes of the depicted genes during reprogramming are shown at the bottom, expressed as transcripts per million (TPM).

## RESULTS

### KLF4 binding during reprogramming induces chromatin opening and precedes enhancer and gene activation

To determine the molecular changes around KLF4 targets during iPSC formation, we first mapped its genome-wide binding at different stages of reprogramming using “reprogrammable” MEFs (Rosa26-M2rtTA/Col1a1-OKSM) induced with doxycycline (dox)^25^ in the presence of ascorbic acid (Fig.1a, bottom). Under these conditions the resulting iPSCs are molecularly and functionally indistinguishable from embryonic stem cells (ESCs) ^26, 27^ and, thus, the term PSCs will be used throughout the text to describe either cell type. For our earliest time point, we collected bulk populations on day 3 after dox treatment, whereas at later stages, on day 6 and day 9, we sorted SSEA1^+^ cells to enrich for cells that are on the right trajectory towards induced pluripotency^14, 25, 28, 29^ (Supplementary Fig.1a). Finally, we used isogenic ESCs and iPSCs^26, 27^ as reference points for established pluripotency. ChIP-seq analysis showed a highly dynamic pattern of KLF4 occupancy during reprogramming with two major categories of binding sites: (i) enriched during intermediate reprogramming stages, but weakly detected in PSCs (Transient KLF4 targets) and (ii) PSC targets, which represent the actual KLF4 binding repertoire once stem cell identity is acquired (Fig.1b and Supplementary Table 1). Among the PSC KLF4 binding sites, 30% were already bound on day 3 (Early KLF4 targets), while the rest were either gradually established during reprogramming (Mid KLF4 targets) or enriched only in established PSCs (Late KLF4 targets). To gain insights into the nature and potential function of each category of KLF4 targets, we performed genomic annotation based on their chromatin state classification introduced by Chronis et al^8^ as well as Gene Ontology (GO) analysis using the GREAT tool^30^ (Fig.1c and Supplementary Fig.1b). Early KLF4 targets mostly enriched for promoters of genes involved in regulation of metabolic processes and cell junction organization, in agreement with the previously reported early role of KLF4 in regulating these processes^14^. On the other hand, Mid and Late KLF4 targets included an increasing number of pluripotency-associated enhancers and enriched for stem cell maintenance genes, including many master regulators of pluripotency, such as *Sox2*, *Nanog*, *Esrrb* and *Klf4*. Finally, Transient KLF4 targets enriched for enhancers previously detected in partially reprogrammed cells^8^ (Supplementary Fig.1b) as well as genes involved in negative regulation of cell cycle, apoptosis and various signaling pathways associated with differentiation, such as TGF-beta signaling (Fig.1c). Therefore, transient KLF4 binding might be associated with unsuccessful reprogramming and alternative fates induced by OKSM expression, as reported in other studies ^31–33^.

The differential kinetics of KLF4 binding prompted us to investigate the epigenetic features of KLF4 targets in MEFs and during reprogramming (Fig.1d, Fig.1e and Supplementary Fig.1c). Integration of ATAC-seq and KLF4 ChIP-seq datasets revealed that ∼60% of the Early KLF4 binding sites were already open in MEFs, suggesting that preexisting chromatin accessibility could partly explain the early binding of KLF4 on these targets (Fig.1d and Fig. 1e). In contrast, the majority (>70%) of Mid and Late KLF4 targets were characterized by closed chromatin configuration in MEFs (Fig.1d and Fig.1e) and higher DNA methylation levels compared to early targets^12^ (Supplementary Fig.1d). These genomic regions gained accessibility concomitantly with KLF4 binding at later timepoints, suggesting the requirement of additional factors for epigenetic remodeling. However, we also observed a large number of inaccessible regions in MEFs that became occupied by KLF4 on day 3 (∼40% of early and ∼75% of transient targets, Fig.1e), indicating that the ability of this TF to access “closed” sites is context-dependent (Fig.1d-e). In agreement, motif enrichment analysis revealed distinct classes of candidate TFs that may synergize with KLF4 earlier or later in reprogramming to promote its stage-specific binding (Supplementary Fig.1e).

KLF4 has been proposed to function both as an activator and repressor of gene expression ^11, 14^. To assess the impact of KLF4 binding on enhancer activity, we performed ChIP-seq for H3K27 acetylation (H3K27ac) in MEFs, PSCs and intermediate reprogramming stages and observed evidence for drastic changes in enhancer usage during iPSC generation (Fig.1f and Supplementary Fig.1f). Less than 5% of decommisioned MEF enhancers (regions that lost H3K27 acetylation between MEFs and day 3) were targeted by KLF4, whereas about 35% of the total acquired PSC enhancers and almost the entirety of (so-called) super enhancers (SE)^34^ were bound by KLF4 concomitantly or prior to H3K27 acetylation (Fig.1g and Supplementary Table 2). Moreover, RNA-seq analysis (Supplementary Fig.1g and Supplementary Table 3) of genes linked to Early, Mid, Late or Transient KLF4 ChIP-seq peaks showed a strong trend for upregulation, rather than downregulation, at the respective stages of reprogramming (Supplementary Fig.1h). These results suggest that KLF4 binding predominantly results in enhancer and gene activation rather than repression during reprogramming. Representative examples for each category of KLF4 target loci are shown in Figure 1h.

In conclusion, our data document stage-specific KLF4 binding with progressive targeting of PSC-associated enhancers, while genes related to “failed” reprogramming trajectories, such as apoptosis or other somatic lineages, were transiently occupied. Globally, the kinetics of KLF4 binding was partly dependent on preexisting chromatin accessibility and DNA methylation levels and either coincided with or preceded enhancer and gene activation.

### Enhancer interactions are extensively rewired between MEFs and PSCs in concordance with epigenetic and transcriptional changes

Previous studies utilizing targeted (4C-seq) or global (HiC) chromatin conformation assays have demostrated that chromatin topology around specific genomic loci and globally at the scale of compartments and Topologically Associated Domains (TADs)^35^, are drastically reorganized during reprogramming^15, 17–21^. However, cell type-specific regulatory loops, such as enhancer-promoter interactions, were under-represented in these studies likely due to technical limitations. Here, we performed H3K27ac HiChIP^36^ in MEFs and PSCs in order to generate high-resolution contact maps around active enhancers and promoters and characterize the degree of architectural reorganization during reprogramming. We called statistically significant interactions at 10kb resolution and within a maximum range of 2MB using Mango^37^ (see Methods) to specifically detect local interactions mediated by H3K27ac. We further refined our set of candidate interactions by considering only loops that overlapped with H3K27ac ChIP-seq peaks in at least one anchor (Supplementary Fig.2a). Differential looping analyses between normalized read-counts (counts-per-million; CPM) of the union of all significant loops called (pvalue<0.1 and Log Fold Change (LogFC) >2 or <-2) revealed about 40,000 contacts that were enriched either in MEFs or in PSCs (Fig.2a and Supplementary Table 4). By applying stringent statistical criteria (pvalue>0.5, logFC<0.5 & logFC>-0.5), we also identified a group of ∼8,000 H3K27ac contacts that show constant interaction strength between MEFs and PSCs. Integration of RNA-seq data showed a significant positive correlation of MEF-specific or PSC-specific H3K27ac loops with increased expression of associated genes in the respective cell type (Fig.2b). These findings demonstrate that H3K27ac HiChIP enables mapping of cell-type specific regulatory contacts and assignment of active enhancers to target genes.

**Figure 2.**
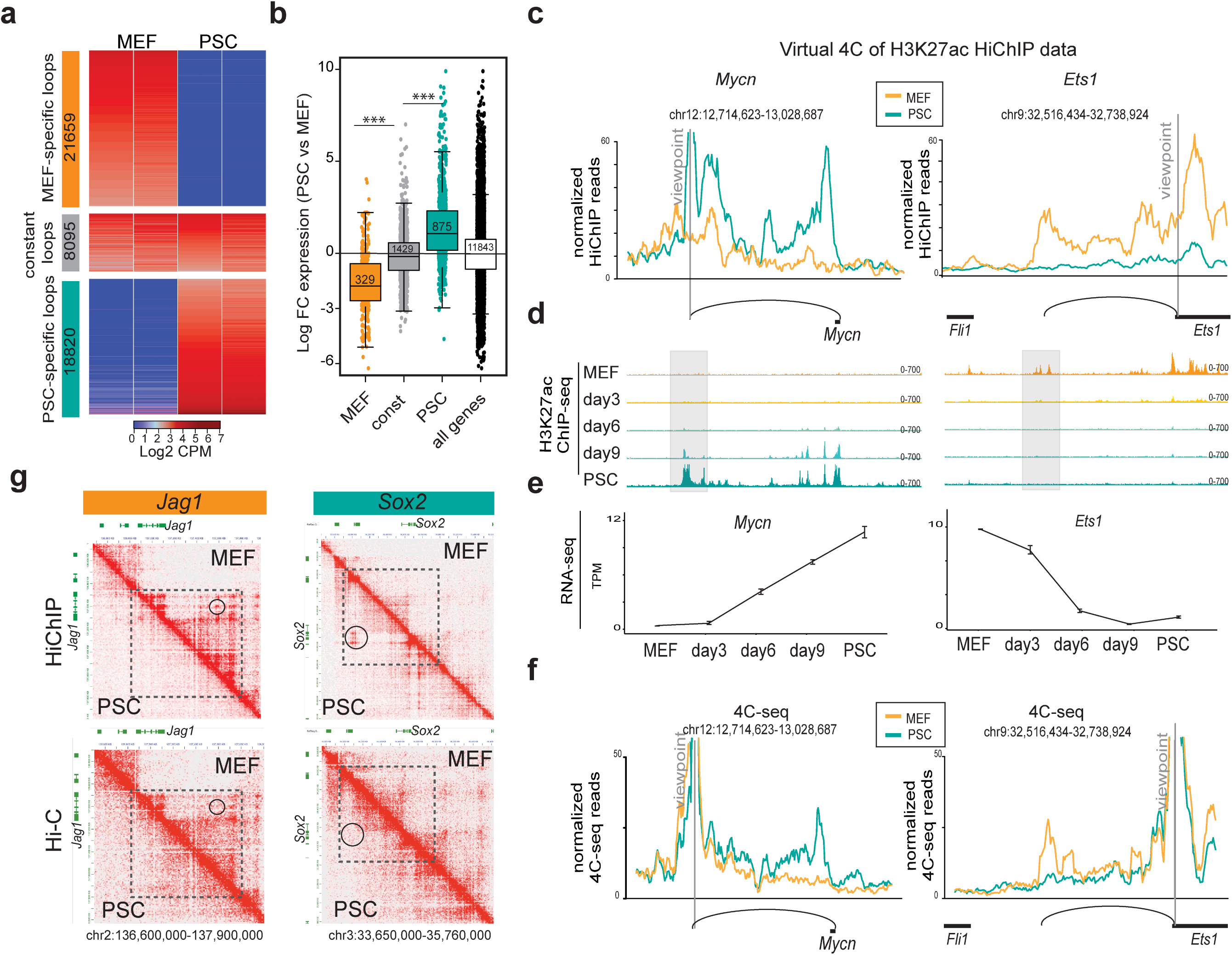
Characterization of 3D enhancer connectomes in MEFs and PSCs by H3K27ac HiChIP analysis. **a**, Heatmap of differential loops detected by H3K27ac HiChIP in MEF versus PSC. Differential loops were called by average logFC>2 or <-2 and p-value < 0.1, constant loops were called by average logFC >-0.5 & logFC <0.5 and p-value >0.5. Heatmap shows Log2 counts-per-million (CPM) per replicate. **b**, RNA expression changes between MEFs and PSCs of genes that were exclusively involved in at least one MEF-specific, PSC-specific or constant H3K27ac loops. All protein-coding genes were used as control. Asterisks indicate significant difference (p<0.001) as calculated by unpaired one-sided t-test. **c**, Virtual 4C representation of normalized H3K27ac HiChIP signals around selected viewpoints (*Mycn* enhancer and *Ets1* promoter). The respective H3K27ac ChIP-seq IGV tracks are shown in **d**, while the RNA changes during reprogramming, expressed as transcripts per million (TPM), are shown in **e**. **f**, 4C-seq analysis around the same viewpoints as in (**c**) validate the presence and cell-type specificity of HiChIP-detected loops. 4C-seq signals normalized by sequencing depth and averaged across replicates are shown. **g**, HiChIP (top) and HiC (bottom) heatmaps generated by Juicebox^76^ at 10Kb resolution around MEF-specific (*Jag1*) or PSC-specific (*Sox2*) contacts. Both PSC and MEF data are shown, separated by the diagonal. Signal indicates CPM normalized per matrix. Dotted squares indicate regions with cell-type specific configuration as detected by both HiC and HiChIP. Circles show examples of cell-type specific contacts that are detected in HiChIP and missed in HiC data.

Examples of cell-type specific enhancer-promoter contacts are shown in Figure 2c, which illustrates normalized H3K27 HiChIP signals in the format of virtual 4C around *Mycn* and *Ets1*. The promoters of these genes establish high-frequency contacts with distal enhancers (>100kb) in a cell-type specific manner. The position and patterns of the detected chromatin loops are in high concordance with acquisition or loss of H3K27ac marks and the respective transcriptional changes during reprogramming (Fig.2d-e). Importantly, high-resolution 4C-seq analysis around *Mycn* enhancer and *Ets1* promoter (Fig.2f) showed a remarkable similarity with the HiChIP results, validating the cell-type specific nature of HiChIP-detected interactions regardless of H3K27ac.

To determine in a global fashion the degree to which differential HiChIP contacts reflect actual chromatin conformation changes, rather than a technical bias due to acquisition or loss of the H3K27ac mark from loop anchors, we performed HiC analysis in MEFs and PSCs. First, we observed that only ∼50% of the HiChIP contacts were also detected in HiC of similar sequencing depth (∼100 million accepted reads per replicate) and using the same loop-calling pipeline (Supplementary Fig.2b). This percentage increased to ∼80% when published ultra-resolution HiC data was used^38^ (∼400 million accepted reads in one replicate, Supplementary Fig.2b), suggesting that sequencing depth is a limiting factor for the ability of HiC to detect HiChIP-enriched loops. Higher local background in HiC might be another limiting factor, as shown by comparing virtual 4C plots of HiChIP and HiC signals around the Tbx3 locus (Supplementary Fig.2c). Examples of contact heatmaps, using HiChIP (Fig. 2g, top) or HiC (Fig. 2g, bottom) data, further illustrate this point: although both depict a cell-type specific configuration around select loci (dotted squares around *Jag1* and *Sox2* genes), there are several cell-type specific loops (circled), which are strongly detected by HiChIP, while are weakly detected or fully absent in HiC. Importantly, when we focused on loops that are detected by both approaches, we observed that MEF-specific or PSC-specific HiChIP loops showed significantly stronger HiC signals in the respective cell type, confirming topological reorganization around these regions (Supplementary Fig.2d). These results, in agreement with previous reports^36, 39^, highlight the increased sensitivity of HiChIP compared to HiC to detect cell-type specific loops.

### Complex 3D connectomes in PSCs are associated with strong enhancer activity

In addition to simple Enhancer-Enhancer, Enhancer-Promoter and Promoter-Promoter interactions, we observed that many genomic regions were involved in more than one loop. The degree of connectivity, as detected by H3K27ac HiChIP, was significantly higher among PSC-specific loops compared to MEF-specific or constant loops, with hundreds of genomic anchors found to be connected with 10 or more (up to 33) distant genes and/or enhancers (Fig.3a and Supplemental Fig. 3a). Analysis of HiC data validated the higher connectivity degree of PSCs compared to MEFs (Supplemental Figure 3b), possibly reflecting the more open and plastic chromatin configuration of this cell type^5, 7, 40^.

**Figure 3.**
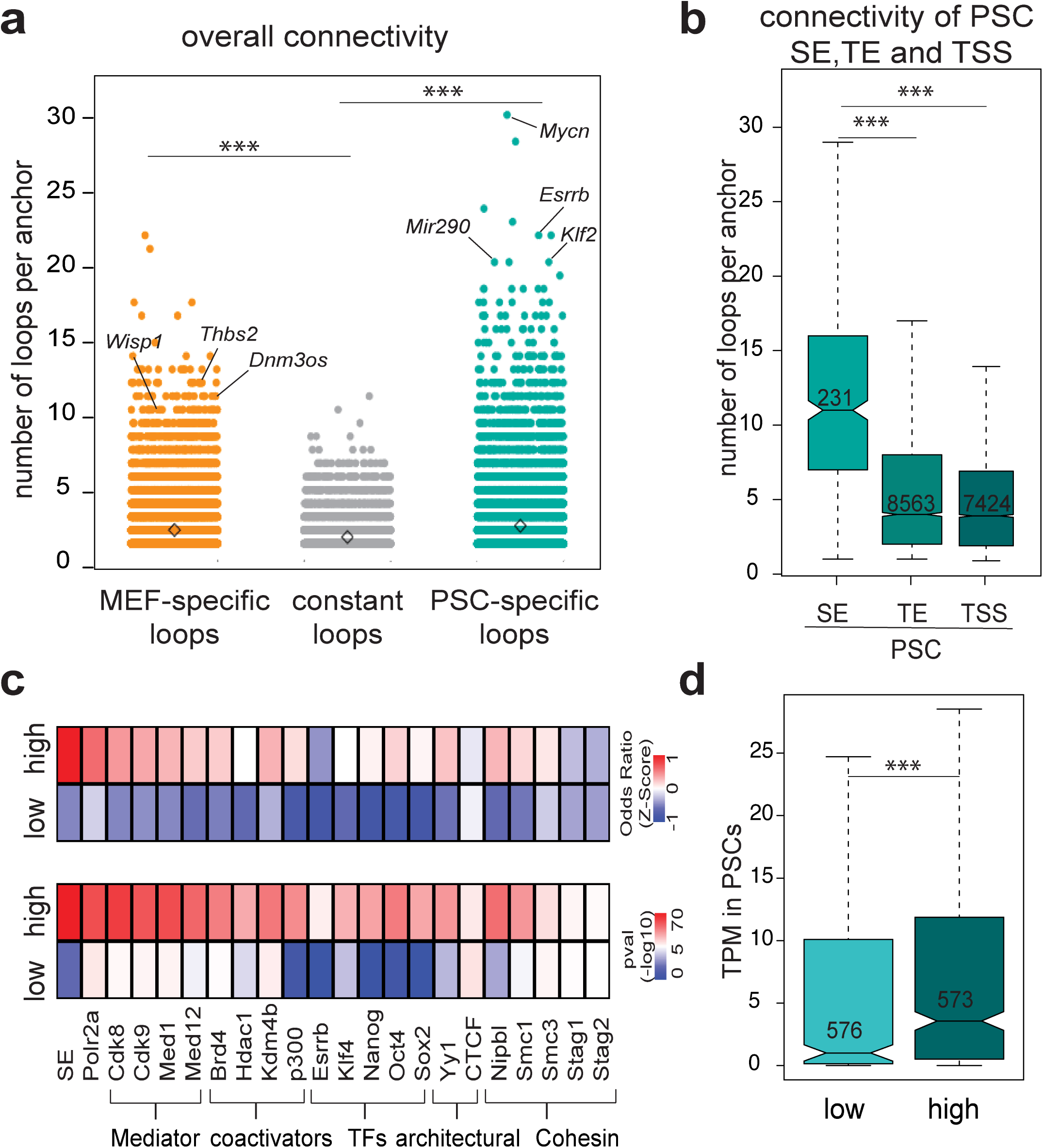
PSC enhancers are characterized by higher connectivity. **a**, Dot plot showing the number of high-confidence contacts (connectivity) around each H3K27ac HiChIP anchor. Asterisks indicate significant difference with p<0.001, as calculated by Wilcoxon rank sum test. **b**, Connectivity of HiChIP anchors containing PSC SE, TE or TSS in PSC. Asterisks indicate significant difference as in (**a) c**, LOLA enrichment analysis of enhancer anchors with low (n=1183) or high connectivity (n=1014) in PSCs using in-house and public ChIP-seq datasets from ESCs (see methods). Heatmaps represent either -log10 p-value (left) or z-score of odds ratio (right). **d**, Expression levels of genes found in low or high connected anchors (expressed in TPM).

Among the highest connected regions in PSCs were critical stem cell regulators, including *Mycn*, *Esrrb*, and *mir290 (*Fig.3a). PSC superenhancers (SE) were also found to be more interactive than typical enhancers (TE)^34^ and transcription start sites (TSS) (Fig.3b). Enrichment analysis of HiChIP anchors based on their connectivity degree (low=1 contact vs high = >4 contacts) showed that highly-connected anchors preferentially associate with binding of Pol II, pluripotency TFs, including KLF4, Mediator complex and transcriptional coactivators (Fig.3c) and connect to highly-transcribed genes (Fig.3d). Cohesin subunits and YY1, which was recently decribed to mediate enhancer-promoter loops^41, 42^, were also preferentially enriched in highly connected anchors, while the classic architectural factor CTCF^43, 44^ was not (Fig.3c). These results suggest that SEs and highly-expressed genes engage in a higher number of chromatin interactions. Importantly, the number of contacts around each enhancer showed poor correlation with the strength of H3K27ac signal (Supplemental Fig.3c), suggesting that our observations are not driven by the biased nature of the HiChIP approach.

### 3D-organized enhancer hubs are associated with coordinated cell-type specific gene expression

To gain insights into the biological role of complex enhancer-promoter interactions, we decided to focus on enhancers that establish connections with multiple gene promoters, potentially forming what we refer to as 3D regulatory hubs (or simply enhancer hubs). Genes found within enhancer hubs were enriched for “stem cell maintenance” categories, including many known pluripotency-associated regulators (e.g. *Zic2*, *Etv2*, *Lin28a*, *Dnmt3l*) (Supplemental Fig.4a) and showed significantly higher expression levels compared to genes with a single-connected enhancer (non-hub genes) or all PSC-expressed genes (Supplemental Fig.4b). Many SE that had been initially assigned to a single gene^34, 45^ were found to either contact individual novel distal target genes or to form hubs with two or more genes of stem cell relevance (e.g. *Utf1*, *Otx2* and *Nacc1*) and high expression levels (Supplemental Fig.4a-b). These results expand the previous pool of candidate genes that are regulated by superenhancers in PSCs^34, 45, 46^. In addition, they raise the possibility that 3D enhancer hubs may coordinate robust expression of stem cell regulatory genes. To test this hypothesis, we selected all protein-coding genes that participate in hubs (2 or more genes contacting the same enhancer) and are differentially expressed between MEFs and PSCs (FC>2, p-adj<0.01) (Supplementary Table 5). We then performed pair-wise comparisons among genes within hubs to calculate the percentage of coregulation (both up- or both down-regulated in PSCs compared to MEFs) or anti-correlation. For control groups we used random gene pairs either of similar linear distance with our test group (global random) or within the same TADs^35^ (TAD-matched random). This approach demonstrated a significant overrepresentation of coregulated gene pairs within enhancer hubs compared to all control groups (Fig.4a) and revealed 311 gene pairs that reside within PSC-enhancer hubs and become concordantly upregulated during reprogramming.

**Figure 4.**
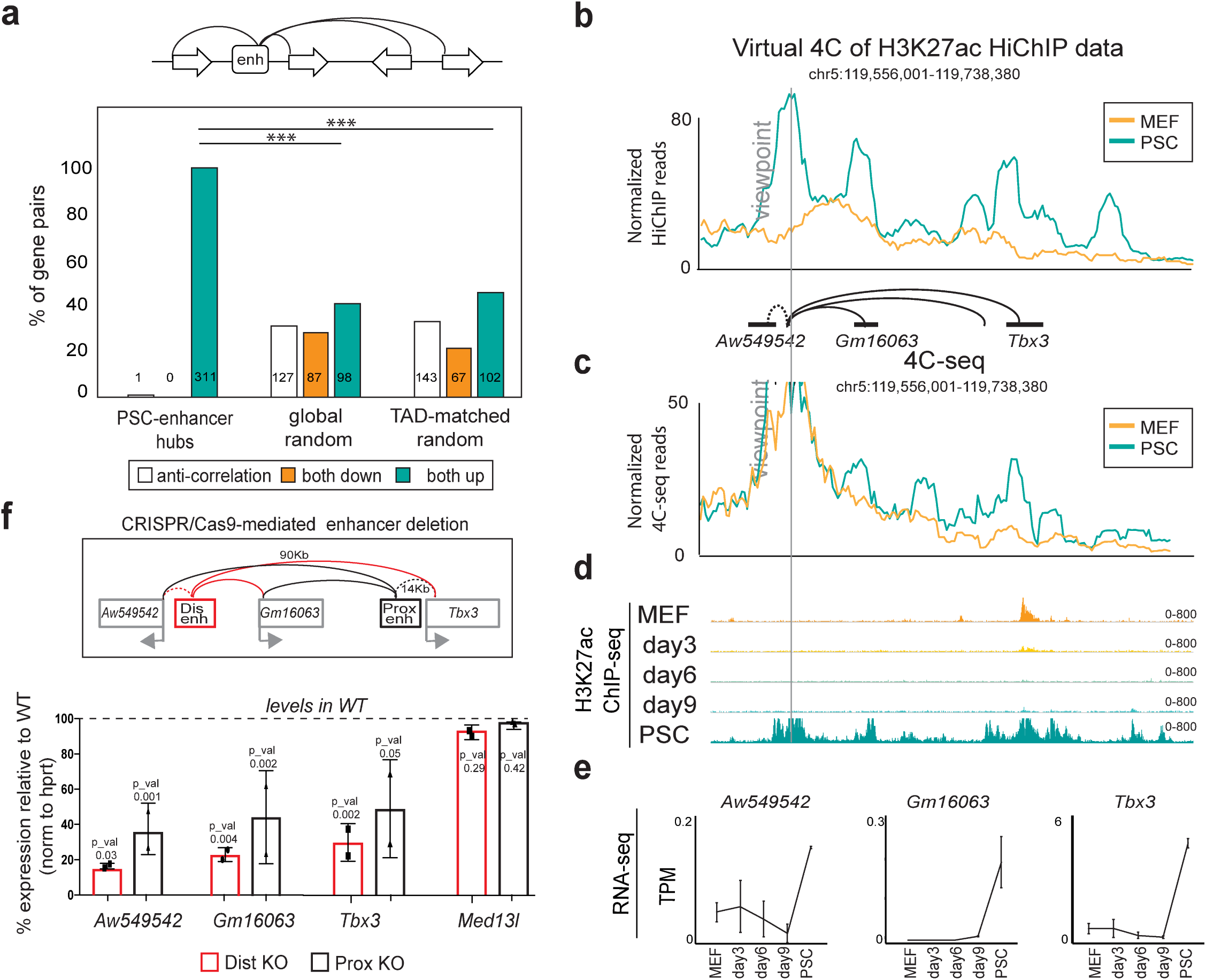
Co-regulation of genes within highly interacting enhancer hubs. **a**, Top: schematic representation of enhancer hubs interacting with two or more gene promoters. Bottom: Barplot indicating the percentage of gene pairs within enhancer hubs that become transcriptionally co-regulated (both up or both down with log2 fold change >=1 or <=-1 & p-adj<=0.01) or anti-regulated (one up and one down) between MEFs and PSCs. Global Random or TAD-matched gene pairs were used as controls (see also Methods). Non-differentially expressed genes were not considered in this analysis (n=487). Significance is indicated by asterisks and was calculated by Fisher’s exact test. **b**, Example of a newly identified enhancer hub in PSCs. Normalized HiChIP signal around the viewpoint is illustrated as a virtual 4C plot. **c**, 4C-seq analysis around the same viewpoint as in (b). 4C-seq counts normalized per sequencing depth are plotted. **d**, H3K27ac ChIP-seq IGV tracks during reprogramming. **e**, RNA-seq signal (TPM) of genes within the hub are shown to highlight coordinated upregulation during reprogramming. **f**, Top: experimental strategy for CRISPR-Cas9-mediated deletions of the *Tbx3* distal (Dis) or proximal (Prox) enhancers within the hub indicated in panel (b). Bottom: RT**-**qPCR showing expression changes of *Tbx3*, *Gm16063*, *Aw54954* and a control gene outside the hub (*Med13l*) in CRISPR-Cas9 engineered PSC carrying homozygous deletions of the distal (Dis-KO) and proximal (Prox-KO) *Tbx3* enhancer calculated as percentage relative to wild=type (WT). P-values are calculated using unpaired one-sided t-test. Error bars indicate standard deviation from n=2 biological replicates. KO: knockout.

To experimentally validate transcriptional coregulation within enhancer hubs, we decided to modulate specific hubs and test transcriptional effects. For this, we focused on an enhancer hub that contacts two proximal non-coding genes (*Aw549542* and *Gm16063)* and the distal (∼90kb) *Tbx3* gene in a PSC-specific manner (Fig.4b). The PSC-specific nature of the HiChIP-detected contacts was validated by 4C-seq (Fig.4c). H3K27ac ChIP-seq and RNA-seq data showed that all connected genes and enhancers within this hub were inactive in MEFs and reprogramming intermediates and became activated only in PSCs, supporting coordinated activation within the hub (Fig.4d-e). Of note, this is not the case for a gene outside the hub (∼800kb), Med13l (Supplemental Fig.4c). Using CRISPR/Cas9 technology^47^ we deleted the distal *Tbx3* enhancer in PSCs, using a deletion of a previously characterized proximal enhancer^48^ *–* which is also part of the same hub – as a reference (Fig. 4f and Supplementary Fig.4d). RT-qPCR analysis of homozygous knock-out (KO) clones demonstrated that the transcriptional levels of *Tbx3* were severely impaired upon disruption of either enhancer (Dis-KO and Pro-KO), with the distal enhancer showing a stronger effect (Fig.4f). Interestingly, the RNA levels of the other hub-connected genes (*Gm1603* and *Aw549542)* were also reduced, while expression of *Med13l* was unaffected (Fig.4f). Furthermore, we used dCas9-KRAB^49^ to target a different enhancer that contacts *Zic2* and *Zic5* genes (Supplemental Fig.4e), which are also coactivated during reprogramming (Supplemental Fig.4f and 4g). CRISPRi-mediated silencing of this enhancer (Supplemental Fig.4h-i) resulted in significant downregulation of both genes, while non-hub genes in linear proximity were only modestly affected (Supplemental Fig.4j).

### KLF4-centered chromatin reorganization during reprogramming associates with enhancer rewiring and transcriptional changes of target genes

Integration of H3K27ac HiChIP results with KLF4 ChIP-seq demonstrated that Early, Mid and Late KLF4 targets (see Fig. 1b) were enriched in PSC-specific H3K27ac interactions, while MEF-specific contacts enriched for transient KLF4 binding (Fig.5a). These results raise the possibility that KLF4 binding is involved in 3D enhancer reorganization during reprogramming. To directly capture the topological changes around KLF4-occupied sites during iPSC formation, we performed KLF4 HiChIP in early (day 3) and mid (day 6) stages of reprogramming as shown in Fig.1a and in PSC. Principle component analysis (PCA) on all statistically-significant interactions called by Mango^37^ distinguished KLF4-bound loops from H3K27ac-marked loops (Supplementary Fig.5a), demonstrating the different nature of chromatin contacts that each antibody captures. Differential looping analysis generated four clusters of dynamic KLF4-centered interactions (Fig. 5b and Supplementary Table 6): two clusters of gained loops, detected either in mid or late reprogramming stages and two clusters of lost loops detected only in early or mid stages.

**Figure 5.**
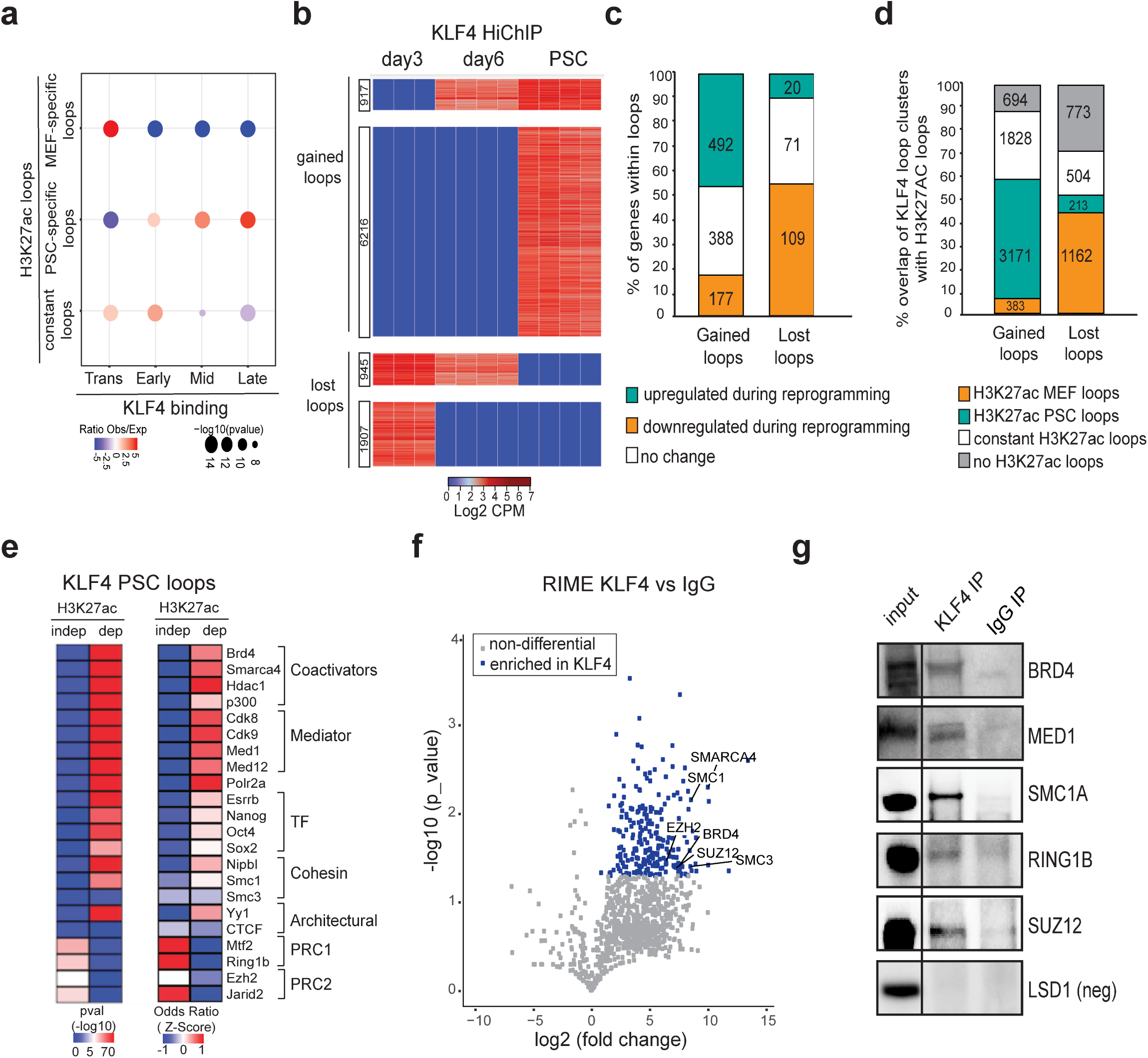
Chromatin reorganization around KLF4 binding sites during reprogramming associates with enhancer rewiring and requires additional cofactors. **a**, Dot plot showing overlap of MEF-specific loops, PSC-specific loops or constant loops as detected by H3K27ac HiChIP with KLF4 Early, Mid, Late and Transient ChIP-seq peaks. The size of the dot represents p-value (as calculated by Fisher’s exact test), while the color indicates the ratio of observed (Obs) versus expected (Exp). **b**, Heatmap of differential KLF4 HiChIP analysis depicting 4 distinct clusters grouped into gained or lost loops. Differential loops were called by average logFC > 2/ or <-2 and p-value < 0.01 between specific pair-wise comparisons (see Methods). Heatmap shows Log2 CPM per replicate. **c**, Stacked barplot indicating the relative proportion of genes within gained or lost KLF4 loops that become upregulated or downregulated (logFC > 1.0, FDR < 0.01 in PSC vs day3) or remain unchanged (logFC > −0.25 & logFC < 0.25) during reprogramming. Numbers of genes per categopry are shown in the respective bars. **d.** Stacked barplot showing the percentage of gained or lost KLF4 loops that were also detected by H3K27ac HiChIP analysis in either MEFs or PSC or in both (constant loops). Note that among all the KLF4 PSC loops, 26% are H3K27ac independent (see Supplementary Figure 4c). **e**, LOLA enrichment analysis of KLF4 binding sites in PSCs that overlap either with H3K27ac-dependent loops (detected by both KLF4 and H3K27ac HiChIP) or -independent (detected only by KLF4 HiChIP). Selected factors that scored as significantly enriched over background are shown. Heatmaps represent either -log10 of p-value (left) or z-score of OddsRatio (right). **f**, Volcano plot showing relative enrichment of proteins that were co-immunoprecipitated with KLF4 versus IgG as identified by RIME (rapid immunoprecipitation mass spectrometry of endogenous protein). Significantly enriched proteins with a p-value< 0.05 and FC >1.5 are colored in blue. Selected co-factors are labeled. **g**, Immunoprecipitation using KLF4 antibody or IgG in PSC extracts followed by western blot analysis validated interaction with selected factors. LSD1 was used as negative control.

To gain insights into the role and nature of the different KLF4-centered loop clusters we investigated the expression changes of associated genes during reprogramming. We found that lost KLF4 HiChIP contacts mostly associate with gene repression, while gained KLF4 loops correlate with gene activation during reprogramming (Fig.5c). Accordingly, comparison with H3K27ac HiChIP data showed that >40% of the lost KLF4 contacts were actually MEF enhancer loops, while >50% of gained KLF4 loops overlapped with PSC enhancer interactions (Fig.5d). Together, these observations support a role of KLF4 binding in the formation/activation of PSC enhancer loops and abrogation/repression of pre-existing somatic loops.

To better understand the relative effect of KLF4 binding and/or looping on gene activation, we focused on enhancer-promoter loops detected by both KLF4 and H3K27ac HiChIP in PSCs and clustered them as: (i) early bound by KLF4 and early formed loops during reprogramming (day 3), (ii) early bound, but late formed loops and (iii) late bound and late formed loops (Supplementary Fig. 5b, left panel). Genes within the first category were robustly upregulated early during reprogramming, while genes in the other two categories were activated only at the late reprogramming stages (Supplementary Fig.5b, right panel). These results indicate that looping coincides with gene activation while KLF4 binding *per se* is not always sufficient to establish promoter-enhancer contacts and activate transcription.

### KLF4 binding engages in both activating and repressive loops in PSCs

Our analysis showed that about 30% of dynamic KLF4-centered loops did not associate with any expression changes and did not overlap with enhancer contacts (Fig.5c,d). Among all KLF4-centered loops in PSCs, 74% overlaps with H3K27ac HiChIP contacts (H3K27ac-dependent), while 26% are H3K27ac-independent (Supplementary Fig.5c). Enrichment analysis using LOLA showed that KLF4 binding sites within H3K27ac-dependent loops are enriched for active enhancer features such as binding of pluripotency TFs (ESRRB, NANOG, SOX2 and POU5F1), YY1 as well as RNA Pol II, co-activators, Cohesin and Mediator subunits (Fig.5e). In contrast, H3K27ac-independent KLF4 anchors are enriched for Polycomb repressive Complex 1 and 2 (PRC1 and PRC2) components, which have been reported to mediate looping among repressed or bivalent genes in PSCs^17, 50, 51^. Genes within H3K27ac-independent KLF4 loops were expressed at significantly lower levels compared to the genes in H3K27ac-dependent loops (Supplemental Fig.5d) and enriched for Gene Ontology categories associated with development and lineage specification (Supplemental Fig.5e). These findings raises the possibility that KLF4 is engaged in chromatin loops with distinct properties and functions, possibly by interacting with different architectural cofactors.

To test the chromatin co-occurrence of KLF4 with computationally-predicted cofactors, we performed RIME^52^ (Rapid Immunoprecipitation Mass spectrometry of Endogenous proteins) in PSCs using either a KLF4 antibody or IgG as control (Fig.5f). This identified 228 high-confidence (FC>1.5 over IgG and p-value<0.05) protein partners (Supplementary Table 7). In addition to novel candidates, RIME detected several of the predicted cofactors, including components of the Cohesin complex, PRC1 and PRC2 as well as co-activators, such as BRD4. Immunoprecipitation using PSC extracts followed by Western blot analysis validated interaction of KLF4 with selected candidates (Fig.5g). These results support the notion that KLF4 participates in different categories of loops in PSCs (Supplemental Fig.5f): (i) activating chromatin loops that are enriched in Cohesin, coactivators and other pluripotency TFs and engage highly-expressed genes involved in cell cycle and stemness (e.g. *Nodal, Mycn, Pou5f1, Dppa2*); (ii) repressive loops mediated by PRC1 and PRC2 components that involve genes related to cell differentiation and development (e.g. *Hoxd10, Bmp4, Serpine3, Fgf9*).

### Depletion of KLF factors in PSCs disrupts a subset of enhancer loops and expression of linked genes

To dissect the role of KLF4 in the 3D enhancer connectome of pluripotent cells, we generated an ESC line that enables dox-inducible targeting of *Klf4* by CRISPR-Cas9. Although KLF4 protein levels were successfully reduced 48 hours after dox addition (Supplementary Fig.6a), we noticed that transcriptional levels of *Klf2* and *Klf5*, encoding TFs with partially redundant function to KLF4^53^, were upregulated in these cells, suggesting compensatory mechanisms (Supplementary Fig.6b). We therefore targeted all three KLF factors using the same conditional system. Shortly after dox induction (24 hours), when the levels of KLF proteins were successfully reduced but before other pluripotency factors such as NANOG were affected (Supplementary Fig.6c), we performed H3K27ac HiChIP and ChIP-seq as well as RNA-seq (Supplementary Table 8). Comparison of enhancer connectomes in uninduced (WT) and induced (triple KO, TKO) cells, revealed 7024 contacts which were consistently reduced (lost) in all TKO replicates and 3488 newly established loops (Fig.6a). The observation that the majority of contacts remained unaffected might be due to residual KLF protein levels (Supplemental Fig.6c) during the intentionally short treatment with dox and/or indicate the presence of additional factors that maintain enhancer architecture and activity. More than 60% of lost loops were bound by KLF4 (ChIP-seq) on one or both anchors, indicating that disruption of these loops is likely a direct effect of KLF factors downregulation (Fig.6b). Of note, multiconnected hubs and superenhancers were preferentially affected compared to typical enhancers, showing a significant reduction in the number of interactions (Supplemental Fig.6d).

**Figure 6.**
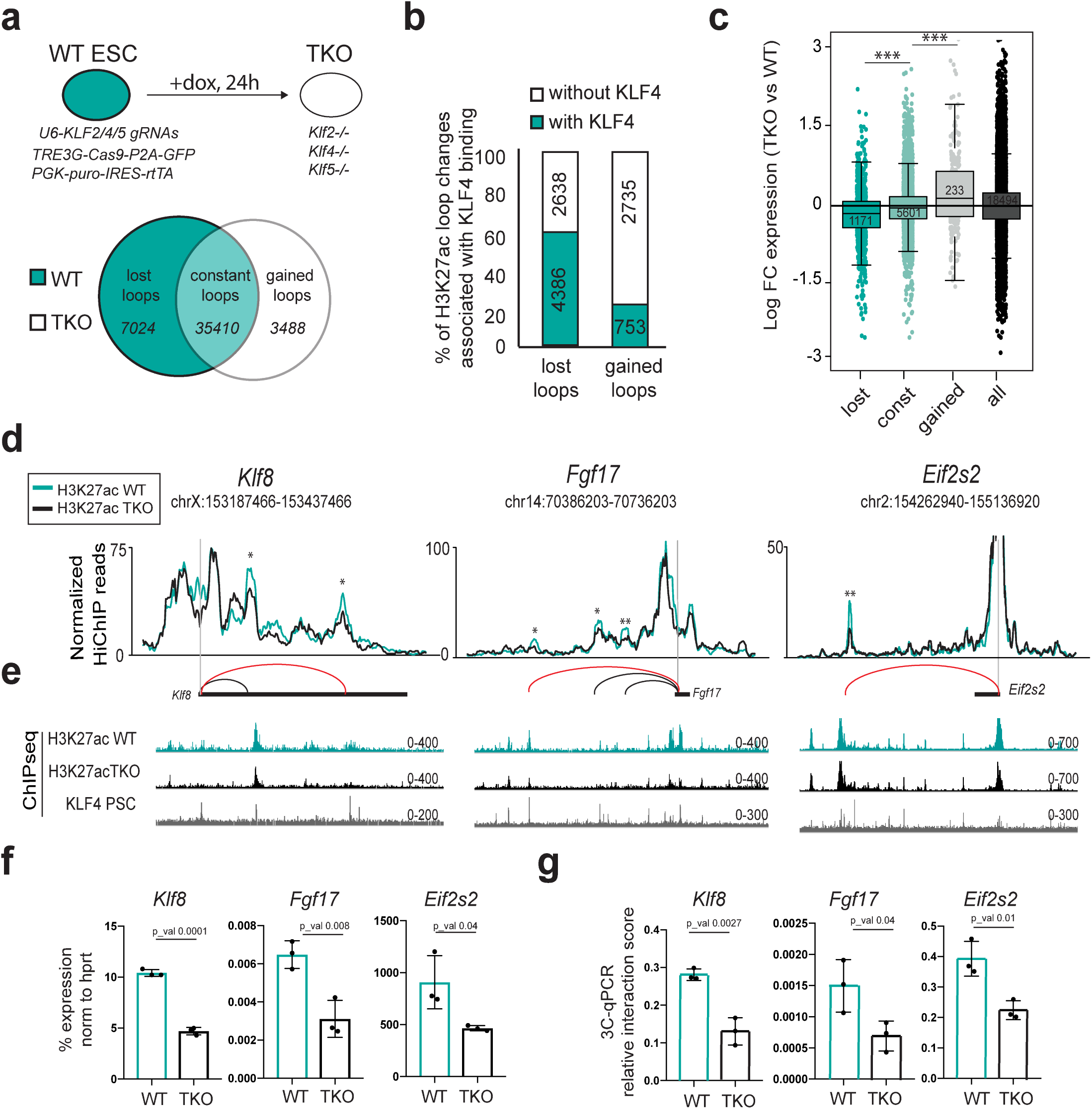
Inducible depletion of KLF proteins induces 3D enhancer reorganization and concordant transcriptional changes. **a**, Top: schematic diagram of the experimental approach used to knock out (KO) KLF2, KLF4 and KLF5 protein in ESCs using a doxycycline (dox)-inducible CRISPR-Cas9 construct. Bottom: Venn diagram showing the number of H3K27ac HiChIP loops that were gained or lost (p-value<0.05 and fold change >1.5 or <-1.5) or remained constant (logFC >-0.5 & <0.5 and p-value>0.5) in triple knock out (TKO) ESCs compared to uninduced (WT) ESCs. **b**, Stacked barplots showing the percentage of gained or lost H3K27ac HiChIP loops in TKO versus WT, whose anchors overlap or not with KLF4 ChIP-seq peaks in PSCs. Numbers represent the actual number of loops. **c**, RNA expression changes of genes within anchors of H3K27ac HiChIP loops (lost, constant or gained loops). All protein-coding genes were used as control. The respective numbers of genes are shown in the boxes. Asterisks indicate significant difference (p<0.001) as calculated by an unpaired one-sided t-test. **d**, Examples of H3K27ac lost loops in TKO vs WT ESC as identified by H3K27ac HiChIP. Normalized H3K27ac HiChIP signals are illustrated in a virtual 4C format around the viewpoints (*Klf8* promoter*, Fgf17* promoter*, Eif2s2* promoter). Asterisks mark the differential loops detected (* p<0.1, ** p<0.01). Statistics were calculated with the R-package edgeR (see Methods for more details). **e**, H3K27ac and KLF4 ChIP-seq tracks around each of the genomic regions indicated in (**d**). **f**, RT**-**qPCR showing expression changes of *Klf8, Fgf17 and Eif2s2* in WT and TKO PSC calculated as percentage relative to *Hprt* levels. P-values were calculated using an unpaired one-sided t-test. Error bars indicate standard deviation from n=3 biological replicates. **g**, 3C-qPCR analysis validating the reduced contact frequency between *Klf8, Fgf17 and Eif2s2* promoters and their respective distal enhancers (marked with a red line in panel (**d)**) in TKO compared to WT ESCs. Unpaired one-sided t-test was used to determine P-values. Error bars indicate standard deviation using n=3 biological replicates.

Integration of RNA-seq data showed that genes within lost or gained loops were significantly down- or up-regulated, respectively, in TKO compared to WT cells (Fig.6c). The relatively moderate transcriptional changes may reflect the short dox-treatment and/or RNA stability. Examples of lost loops, represented as a virtual 4C of H3K27ac HiChIP data in WT and TKO cells, along with the respective KLF4 and H3K27ac ChIP-seq tracks are shown in Figure 6d and 6e. The reduced mRNA levels of *Klf8*, *Fgf17* and *Eif2s2* genes and the disruption of the respective gene-enhancer contacts in TKO cells were independently validated by RT-qPCR and 3C-qPCR, respectively (Fig.6f and 6g). These results demonstrate that depletion of KLF factors in PSCs results in abrogation of thousands of enhancer contacts genome-wide and concordant dysregulation of connected genes.

### Disruption of KLF4 binding sites interferes with enhancer looping and transcriptional activation

To ascertain whether KLF4 binding is critical for maintenance of 3D enhancer contacts in PSCs, we targeted KLF4 binding sites within selected enhancer hubs and examined local topological and transcriptional effects. We initally chose the distal *Tbx3* enhancer, deletion of which resulted in downregulation of all three hub-connected genes (Fig.4f). The multiple contacts of this enhancer with the surrounding genes were detected both by H3K27ac and KLF4 HiChIP only in PSCs but not in MEFs or reprogramming intermediates (Fig.7a). This is in concordance with the late binding of KLF4 to this enhancer (Fig.7b) and the late transcriptional activation of the entire locus (Fig.4e). We utilized CRISPR/Cas9 technology to disrupt the strongest KLF4 binding motif within this enhancer hub (Fig.7c and Supplemental Fig.7a-c). Four different homozygous mutant clones were validated for impaired KLF4 binding by ChIP-qPCR (Supplemental Fig.7d) and used for further characterization. RT-qPCR analysis demostrated that the transcriptional levels of all hub-connected genes *(Aw549542, Gm1603* and *Tbx3*) were significantly reduced, whereas the expression of a gene outside the hub was not affected (*Med13l*) (Fig.7d). Consistent with transcriptional downregulation, the long-range contacts between the enhancer hub and its target genes were significantly weakened in mutant clones as shown by 3C-qPCR (Fig.7e), while the interaction of *Tbx3* with the proximal enhancer or a KLF4-independent contact in a different genomic region remained unaffected (Fig.7e). Using a similar approach, we mutated a strong KLF4 binding site within the previously described *Zfp42* superenhancer^54^, which contacts both *Zfp42* and the distal (∼150kb) *Triml2* gene in a PSC specific manner (Fig.7f-g). Homozygous mutant ESCs showed significant downregulation of *Zfp42* expression and a concordant reduction of enhancer-*Zfp42* promoter contact frequency (Fig.h-j). Intringuingly, the expression levels of *Triml2* remained unaffected in the mutant clones and the connection with the enhancer appeared even stronger (Fig.7i-j), suggesting that KLF-dependent and independent mechanisms may regulate looping and activity of the same enhancer on different genes. Taken together, these results provide evidence for a dual role of KLF4 as a transcriptional regulator and chromatin organizer in PSCs.

**Figure 7.**
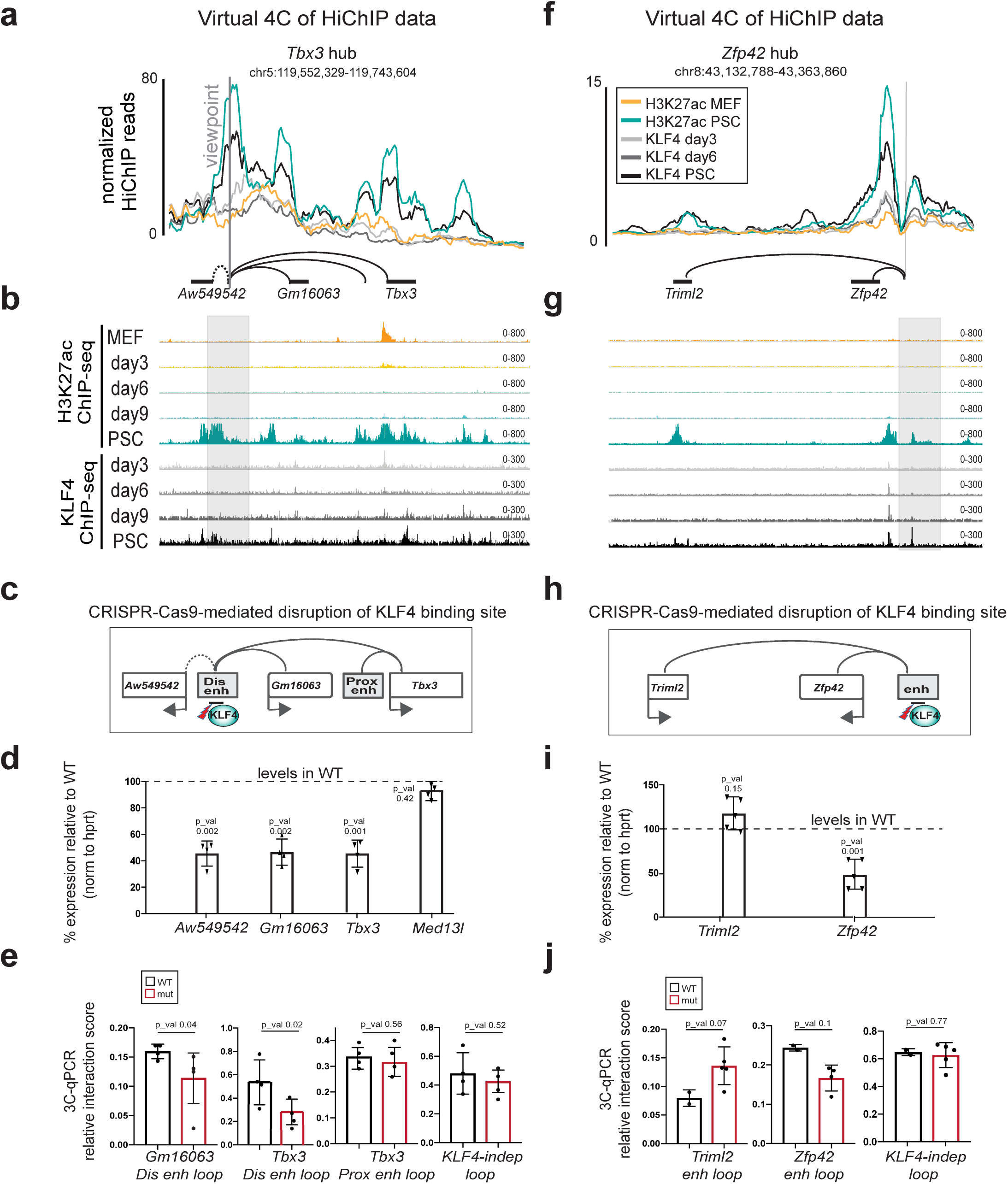
Disruption of KLF4 binding site within *Tbx3* and *Zfp42* enhancers induces looping abrogation and downregulation of linked genes in PSCs. **a**, Normalized KLF4 and H3K27ac HiChIP signals are illustrated as virtual 4C line plots around the *Tbx3* distal enhancer hub (see also Fig.4b-f). The respective ChIP-seq IGV tracks are shown in **b**. **c**, Schematic illustration of the CRISPR-Cas9 targeting strategy to generate mutated KLF4 binding motifs (mut) within the distal *Tbx3* enhancer. **d**, RT**-**qPCR showing expression changes of hub-associated genes (*Tbx3*, *Gm16063* and *Aw54954)*. *Med13l* is used as control gene outside the hub. Values were calculated as percentage relative to wild type (WT) after normalization relative to *Hprt* mRNA levels. Unpaired one-sided t-test was used to determine significance relative to WT (p-value is indicated on the top of each bar). Error bars indicate standard deviation from n=4 different PSC clones carrying homozygous mutations of KLF4 binding motif (mut). **e**, 3C-qPCR analysis showing the relative interaction frequency of *Tbx3* distal enhancer with the promoters of linked genes in WT and mutant (mut) clones. Error bars indicate standard deviation. *n=*2 for WT and n=4 for mut biological replicates. Unpaired one-tailed Student’s t-test was used to determine significance relative to WT (the value is indicated on the top of each bar). **f-j**, Representation, analysis and functional validation of *Zfp42* enhancer hub similarly to panels (a-e) for *Tbx3* hub. The same normalizations and statistical tests were applied, with the only difference that n=5 mutant clones carrying homozygous mutations of KLF4 binding motif were used.

## DISCUSSION

Here, we describe the genome-wide dynamics of KLF4 binding and probe its effects on chromatin accessibility, enhancer activity, gene expression and 3D enhancer organization during iPSC reprogramming and in established PSCs. Our data suggest that the kinetics of KLF4 binding and the temporal relationship with gene and enhancer activity is partly dependent on preexisting chromatin accessibility, the presence of epigenetic barriers such as DNA methylation and/or the availability of additional TFs and cofactors, such as ESRRB or NANOG^8, 12, 13^. Nevertheless, KLF4 also binds to chromatin regions that are inaccessible and highly methylated in somatic cells, which is in agreement with its documented ability to act as a pioneer factor and induce chromatin opening and DNA demethylation^9, 55, 56^ and/or its cooperative binding with other reprogramming TFs^8^.

Previous studies utilizing 4C or HiC have characterized dynamic 3D architectural changes during reprogramming either at a small-scale, around specific genomic sites^15, 18^, or at a large-scale, mostly at the levels of compartments and domains^21^. These studies offered important insights into the principles of topological reorganization during cell fate transitions, but they did not capture the dynamic assembly and disassembly of cell-type enhancer contacts. Here, we chose to apply H3K27ac HiChIP analysis, which was reported to have significantly higher discovery rate for cell-type specific loops compared to HiC and Capture HiC methods^36, 39^. Indeed, our data revealed dramatically rewired enhancer connectomes between MEFs and PSCs generating a reference map of cell-type specific regulatory loops. Independent 4C-seq and HiC experiments largely validated the cell-type specific nature of the detected HiChIP interactions, but also revealed technical biases and limitations for each approach, highlighting the need for a deeper and systematic comparison of different 3C assays and analytical tools. Our H3K27ac HiChIP analysis uncovered a set of highly-connected enhancers, which communicate with strongly expressed cell-type specific genes, supporting that high interactivity might be an inherent characteristic of critical regulatory elements for cell identity, as it has been suggested in previous studies^57, 58^. Moreover, we identified a number of cell-type specific enhancers, including many SE, which frequently interact with two or more coregulated genes, supporting a potential role for such hubs in coordinating target gene activation, as previously shown in different contexts^59^. In further support, deletion or inactivation of enhancer hubs resulted in coordinated downregulation of all connected genes without affecting neighboring non-hub genes. Recently developed technologies that capture multiway interactions^60–63^ will enable dissecting to what extent these enhancer hubs represent multiple contacts occuring in the same cell and allele or highly dynamic contacts with one gene at a time. In either case, our results provide genome-wide evidence for the role of selected enhancers in coordinating gene regulation during acquisition and maintenance of pluripotency and demonstrate the potential of this approach to identify novel candidate genes and enhancers critical for specific cellular identities.

There is increasing evidence that TFs are involved in mediating chromatin contacts in different cellular contexts^21, 38, 39, 41, 42, 64–69^, although the underlying mechanisms and the temporal relationships between TF binding and topological and transcriptional changes remain elusive. Encouraged by previous studies reporting potential architectural functions for various KLF protein members^18, 23, 24^, we went on to capture for the first time in a direct and genome-wide manner the dynamic chromatin reorganization around KLF4-binding sites during iPSC formation by KLF4 HiChIP. This approach revealed that KLF4 binding associated with *de novo* establishment of enhancer loops during reprogramming, promoting transcriptional upregulation of linked genes. We also observed that KLF4 binding was not always sufficient for looping formation and gene activation, suggesting the requirement of additional architectural factors and coregulators. In support of this notion, our computational and proteomics analyses revealed distinct sets of candidate cofactors that interact with KLF4 protein either in the context of activating enhancer loops or repressive/poised loops in PSCs. How these proteins work together to form 3D chromatin contacts remains to be shown. Recruitment of architectural cofactors capable to physically tether distal DNA elements is a plausible scenario and is supported by the fact that KLF4 directly interacts with cohesin subunits^43^. Another possibility is that formation of activating or repressive topological assemblies, such as 3D enhancer hubs or polycomb bodies^17, 50, 57, 62, 70, 71^, is the result of “self-organization” through multiprotein condensation. In support of this model, KLF4 and validated cofactors, such as Mediator and BRD4, are charaterized by extensive intrinsically disordered regions (IDRs), which have been shown to promote multivalent interactions and formation of subnuclear condensates^72–75^.

In contrast with previous studies that described the involvement of KLF4 in the maintenance of selected chromatin loops^18, 22^, our study provides evidence for a functional role in the organization and regulation of 3D enhancer contacts and hubs in PSCs at a genome-wide scale. In addition to the global topological effects induced by KLF protein depletion, we showed that targeting individual KLF4 binding sites within specific enhancer hubs was -in some cases- sufficient to disrupt enhancer-promoter contacts and induce downregulation of associated genes. Systematic functional interrogation of KLF4-bound enhancer hubs as identified by HiChIP may enable a deeper understanding of KLF4-dependent and independent mechanisms of topological organization and the establishment of new criteria for identification and functional prioritization of critical regulatory nodes for PSC identity.

## ACKNOWLEDGEMENTS

We are grateful to Ari Melnick and members of the Apostolou, Tsirigos and Stadtfeld lab for critical reading of the manuscript. We also want to thank Luke Dow for sharing the CRISPR/Cas9 vectors. DCG was supported by the New York Stem Cell Foundation and the Family-Friendly Postdoctoral Initiative at Weill Cornell Medicine. AA is supported by a Medical Scientist Training Program grant from the National Institute of General Medical Sciences of the National Institutes of Health (NIH) under award number T32GM007739 to the Weill Cornell/Rockefeller/Sloan Kettering Tri-Institutional MD-PhD Program. This work was funded by the NIH Director’s New Innovator Award (DP2DA043813) and the Tri-Institutional Stem Cell Initiative by the Starr Foundation.

## AUTHOR CONTRIBUTIONS

EA conceived, designed, supervised the study and wrote the manuscript together with DCDG with help from all authors. DCDG performed all experiments with help from DK and VS. AK and APP performed all HiChIP and HiC analyses and integrative computational analysis under the guidance of AT. YL performed initial ChIP-seq, RNA-seq and ATAC-seq analysis. DM performed HiC and CRISPRi experiments using a stable dCas9-KRAB ESC line generated by BA. AA performed RIME experiments and iPSC ChIP-seq. PC and ND run and analyzed RIME results. MS provided reprogrammable cells and guidance on reprogramming experiments.

## DATA AVAILABILITY STATEMENT

All data (RNA-seq, ChIP-seq, ATAC-seq and HiChIP) were submitted to GEO under the accession code GSE113431.

## METHODS

### Cell lines, culture conditions and reprogramming experiments

Mouse ES V6.5 were cultured on irradiated feeder cells in KO-DMEM media (Invitrogen) supplemented with 15% heat-inactivated fetal bovine serum, GlutaMAX, penicillin-streptomycin, non-essential amino acids, β-mercaptoethanol and 1000 U/ml LIF, with or without the presence of 2i (1uM MEKinhibitor (Stemgent 04-0006) and 3uM GSK3 inhibitor (Stemgent 04-0004)).

Mouse embryonic fibroblasts (MEFs) were isolated from a “reprogrammable” mouse harboring a polycystronic OKSM cassette in the *Col1a1* locus and M2rtTA in the *Rosa26* locus^25^. Cells were reprogrammed in the presence of 1ug/ml doxycycline and 50ug/ml ascorbic acid and cultured in ES medium as described above. Cells were collected at the indicated time points.

### Lentiviral production and infection

293T cells were transfected with overexpression constructs along with packaging vectors VSV-G and Delta8.9 using PEI reagent (PEI MAX, Polyscience #24765-2). Supernatant was collected after 48hrs and 72hrs and virus was concentrated using Polyethylglycol (PEG, Sigma # P4338). V6.5 cells were infected in medium containing 5ug/ml polybrene (Millipore, TR-1003-G) followed by centrifugation at 2100rpm for 90 mins at 32°C.

### MACS and FACS

For isolating the SSEA1 positive cells from reprogramming intermediates at day6 and day9 we used magnetic microbeads conjugated to anti-SSEA1 antibody (MACS Miltenyi Biotec #130-094-530) as per manufacture instructions. SSEA positive and negative fractions were then stained for FACS analysis with an anti-Thy1 antibody conjugated to pacific blue fluorophore (ebioscience # 48-0902-82) and anti-SSEA antibody conjugated to APC fluorophore (biolegend #125608).

### Generation, selection and validation of KO cell lines

gRNAs were cloned into the px458 vector (Addgene #48138) using the BbsI restriction enzyme. 0.3 million ESC cells (V6.5) were transfected using 2ug of Left-*Tbx3*-plasmid and 2ug of Right-Tbx3-plasmid (for *Tbx3* enhancer deletions) or 4ug of *Tbx3*-KLF4mut-vector (mutation of KLF4 binding site within *Tbx3* distal enhancer) or 4ug KLF4-Zfp42mut (mutation of KLF4 binding site within *Zfp42* enhancer). DNA was pre-mixed with 50ul media with no additions, and in a separate tube 10ul of Lipofectamine 2000 (Invitrogen #11668019) was pre-mixed with 50ul media with no addition. After 5 minutes the two tubes were combined and incubated at room temperature for 20 more minutes. Cells were then added to the solution and plated on a gelatinized 12 well plate. 48hrs post-transfection, GFP positive single cells were sorted by FACS into 96 well plates. Genotyping of the single cell colonies was performed using a three-primer strategy (for deletions) or by surveyor with T7 (for in-del mutation). Four (*Tbx3* hub) or five (*Zfp42* hub) colonies with homozygous mutations (or w.t. colonies as control) were expanded and used for RT-qPCR and 3C experiments. All gRNA, 3C and RT-qPCR primers are described in Supplementary Table 9.

### CRISPRi of Zic2/5 enhancer

V6.5 cells were infected with lentiviruses harboring the pHR-SFFV-dCas9-BFP-KRAB vector (Addgene,46911) in which the SFFV promoter was replaced with an Ef1a promoter. BFP expressing cells were selected by three rounds of FACS sorting. The resulting V6.5, stably expressing the KRAB-dCas9, were then infected with a lentivirus harboring the pLKO5.GRNA.EFS.PAC vector (Addgene, 57825) containing two gRNAs targeting the Zic2/5 enhancer. Cells were selected with Puromycin (LifeTech K210015) for two days and subsequently collected for RT-qPCR. gRNA and RT-qPCR primers are described in Supplementary Table 9.

### Generation of TKO cell line

V6.5 cells were infected using lentiviruses harboring the c3GIC9 vector^77^ (TRE3G-Cas9-P2A-GFP-PGK-Puro-IRES-rtta) containing gRNA/s targeting either *KLF4* only or *KLF2, KLF4* and *KLF5* in tandem. Following infection cells were selected using Puromycin (LifeTech K210015) and clonal populations were manually picked. Expression of CRISPR-Cas9 from these stable cell lines was induced by addition of Doxycycline for 72hrs (1:1000 dilution of 2mg/ml stock) and KO efficiency in each clonal population was verified by WB: KLF4 (R&D, AF3158) KLF5 (R&D AF3758) KLF2 (Novus biologicals, NBP6181) ESRRB (PPMX, PPH6705) NANOG (Bethyl laboratories, A300-397A) ACTIN (abcam, ab49900). Successful KO clones were then used for subsequent experiments (qPCR, ChIP-seq, 3C and HiChIP) after induction with doxycycline for the indicated time points.

### 3C-qPCR

For each sample 1 to 2 million cells were lysed in 300ul of lysis buffer (10mM Tris-HCl pH 8.0, 10mM NaCl, 0.2% Igepal CA630 with protease inhibitors) and incubated on ice for 20 mins. Cells were centrifuged 2500g for 5min at 4°C and pellet washed once in lysis buffer. Pellets were resuspended in 50ul of 0.5% SDS and incubated at 65°C for 10 mins. 145ul of water and 25ul of 10% triton were added to the samples and incubated 15mins at 37°C. 100 Units of MboI restriction enzyme and 25ul of NEB buffer 2 were added and incubated over night at 37°C with rotation. Next day the enzyme was inactivated at 65°C for 20 mins. The ligation reaction was carried out over night at 16°C by adding 120ul of NEB T4 ligase buffer, 100ul of 10% Triton, 6ul of 20mg/l BSA, 100ul of 10mM ATP and 5ul of T4 ligase (NEB #M0202). The following day, 50ul of 20mg/ml proteinase K and 120ul of 10% SDS were added and the samples were incubated over night at 65°C. Lastly, 10ul of 10mg/ml RNAse was added and samples incubated 1 hour at 37°C. Following phenol chloroform purification, the DNA was precipitated using 1.6 volumes of 100% ethanol and 0.1 volume of 3M sodium acetate. After incubation at −80°C for 1 hour samples were spun for 15mins at 4°C at 16000rpm. Pellets were washed twice with 70% ethanol and dissolved in 100ul of 10mM Tris pH8. Qbit was used to measure sample concentrations and 100ng of material was used to amplify the desired regions by qPCR. All primer sequences can be found in Supplementary table 9.

### ChIP-seq

ChIP-seq was performed as previously described^78^. Specifically, cells were crosslinked in 1% formaldehyde at RT for 10 minutes and quenched with 125mM glycine for 5 mins at RT. 50 million cells were used for KLF4 ChIPs and 10 million for H3K27acetylation ChIP. Cell pellets were washed twice in PBS and resuspended in 400ul lysis buffer (10mM Tris pH8, 1mM EDTA, 0.5% SDS) per 20 million cells. Cells were sonicated in a bioruptor device (30 cycles 30sec on/off, high setting) and spun down 10 minutes at 4°C at maximum speed. Supernatants were diluted 5 times with dilution buffer (0.01%SDS, 1.1% triton,1.2mM EDTA,16.7mM Tris pH8, 167mM NaCl) and incubated with the respective antibody (2-3ug/10M cells) (KLF4 R&D #3158, H3K27ac ab4729) O/N with rotation at 4°C. Next day, protein G Dynabeads (ThermoScientific) preblocked with BSA protein (100ng per 10ul Dynabeads) were added (10ul blocked Dynabeads per 10 million cells) and incubated for 2-3 hours at 4°C. Beads were immobilized on a magnet and washed twice in low salt buffer (0.1% SDS,1% triton, 2mM EDTA, 150mM NaCl, 20mM Tris pH8), twice in high salt buffer (0.1% SDS,1% triton, 2mM EDTA, 500mM NaCl, 20mM Tris pH8), twice in LiCl buffer (0.25M LiCl, 1% NP40, 1% deoxycholic acid (sodium salt), 1mM EDTA, 10mM Tris pH8) and once in TE. DNA was then eluted from the beads by incubating with 150ul elution buffer (1% SDS, 100mM NaHCO3) for 20 minutes at 65°C (vortexing every 10min). Supernatants were collected and reverse-crosslinked by incubation at 65°C O/N in presence of proteinase K. After RNase A treatment for 1hr at 37°C, DNA was purified using the minElute kit (Qiagen). 6-10ng of immunoprecipitated material was used for ChIP-seq library preparation using the KAPA Hyper prep kit (KAPA Biosystems). Libraries were sequenced on an Illumina HiSeq 2500 platform on SE50 mode.

### ATAC-seq

ATAC-seq was performed as previously described^79^. In brief, a total of 50,000 cells were washed once with 50 µL of cold PBS and resuspended in 50 µL lysis buffer (10 mM Tris-HCl pH 7.4, 10 mM NaCl, 3 mM MgCl2, 0.2% (v/v) IGEPAL CA-630). The suspension of nuclei was then centrifuged for 10 min at 800 g at 4°C, followed by the addition of 50 µL transposition reaction mix (25 µL TD buffer, 2.5 µL Tn5 transposase and 22.5 µL nuclease-free H_2_O) using reagents from the Nextera DNA library Preparation Kit (Illumina #FC-121-103). Samples were then incubated at 37°C for 30min. DNA was isolated using a ZYMO Kit (#D4014). ATAC-seq libraries were first subjected to 5 cycles of pre-amplification. To determine the suitable number of cycles required for the second round of PCR the library was assessed by quantitative PCR as described in Buenrostro et al ^79^ and the library was then PCR amplified for the appropriate number of cycles using Nextera primers. Samples were subject to a dual size selection (0.55x-1.5x) using SPRI beads (Beckman Coulter #B23317). Finally, the ATAC libraries were sequenced on a HiSeq 2500 platform on PE50 mode.

### RNA-seq

Total RNA was prepared with TRIZOL (Life technologies #15596018) following manufacturer’s instructions. Libraries were generated by the Weill Cornell Genomics core facility using the Illumina TruSeq stranded mRNA library preparation kit (#20020594) and sequenced on an Illumina HiSeq4000 platform on SE50 mode.

### HiChIP

HiChIPs were performed as previously described^36^ with some modifications. In brief, up to 15 million crosslinked cells (for KLF4 HiChIPs two samples of 15 million cells were combined at the end, for each sample replicate) were resuspended in 500 µL of ice-cold HiC lysis buffer (10 mM Tris-HCl pH 7.5, 10 mM NaCl, 0.2% NP-40, 1x protease inhibitors) and rotated at 4°C for 30 min. Nuclei were pelleted and washed once with 500 µL of ice-cold HiC lysis buffer. Pellet was then resuspended in 100 µL of 0.5% SDS and incubated at 62°C for 10 min. 285 µL of water and 50 µL of 10% Triton X-100 were added, and samples were rotated at 37°C for 15 min. 50 µL of NEB Buffer 2 and 15 µL of 25 U/µL MboI restriction enzyme (NEB, R0147) were then added, and sample was rotated at 37°C for 2 h. MboI was then heat inactivated at 62°C for 20 min. We added 52 µL of incorporation master mix: 37.5 µL of 0.4 mM biotin–dATP (Thermo Fisher, 19524016); 4.5 µL of a dCTP, dGTP, and dTTP mix at 10 mM each; and 10 µL of 5 U/µL DNA Polymerase I, Large (Klenow) Fragment (NEB, M0210). The reactions were then rotated at 37°C for 1 h. 948 µL of ligation master mix was then added: 150 µL of 10x NEB T4 DNA ligase buffer with 10 mM ATP (NEB, B0202), 125 µL of 10% Triton X-100, 3 µL of 50 mg/mL BSA (Thermo Fisher, AM2616), 10 µL of 400 U/µL T4 DNA Ligase (NEB, M0202), and 660 µL of water. The reactions were then rotated at room temperature for 4 h. After proximity ligation, the nuclei were pelleted and the supernatant was removed. The nuclear pellet was brought up to 880 µL in Nuclear Lysis Buffer (50 mM Tris-HCl pH 7.5, 10 mM EDTA, 0.5% SDS, 1x Roche protease inhibitors, 11697498001), and sonicated with a Bioruptor 300 (Diagenode) for 8 cycles of 30sec each, on a medium setting. Clarified samples were transferred to Eppendorf tubes and diluted five times with ChIP Dilution Buffer (0.01% SDS, 1.1% Triton X-100, 1.2 mM EDTA, 16.7 mM Tris-HCl pH 7.5, 167 mM NaCl). Cells were precleared with 30 µL of Protein G dynabeads (Life technology #10004D) in rotation at 4°C for 1 h. Supernatants were transferred into fresh tubes and antibody was added (8 µg of KLF4 antibody or 3ug H3K27Ac antibody for 15 million cells) and incubated overnight at 4°C. The next day 30 µL of Protein G dynabeads were added to samples and rotated at 4°C for 2 h. After bead capture, beads were washed three times each with low-salt wash buffer (0.1% SDS, 1% Triton X-100, 2 mM EDTA, 20 mM Tris-HCl pH 7.5, 150 mM NaCl), high-salt wash buffer (0.1% SDS, 1% Triton X-100, 2 mM EDTA, 20 mM Tris-HCl pH 7.5, 500 mM NaCl), and LiCl wash buffer (10 mM Tris-HCl pH 7.5, 250 mM LiCl, 1% NP-40, 1% sodium deoxycholate, 1 mM EDTA, make fresh). Samples were eluted with 150 µL of DNA elution buffer (50 mM sodium bicarbonate pH 8.0, 1% SDS, freshly made) and incubated at 37°C for 30 min with rotation. Supernatant was transferred to a fresh tube and elution repeated with another 150 µL elution buffer. 5 µL of Proteinase K (20mg/ml) (Thermo Fisher) were added to the 300 µL reaction and samples were incubated overnight at 65°C. Samples were purified with DNA Clean and Concentrator columns (Zymo Research) and eluted in 10 µL of water. Post-ChIP DNA was quantified by Qubit (Thermo Fisher) to estimate the amount of Tn5 (Illumina) needed to generate libraries at the correct size distribution (see below). 5 µL of Streptavidin C-1 beads (Thermo Fisher) were washed with Tween Wash Buffer (5 mM Tris-HCl pH 7.5, 0.5 mM EDTA, 1 M NaCl, 0.05% Tween-20) then resuspended in 10 µL of 2x biotin binding buffer (10 mM Tris-HCl pH 7.5, 1 mM EDTA, 2 M NaCl). Beads were added to the samples and incubated at room temperature for 15 min with shaking. After capture, beads were washed twice by adding 500 µL of Tween Wash Buffer and incubated at 55°C for 2 min with shaking. Samples were then washed in 100 µL of 1x TD Buffer (2x TD Buffer is 20 mM Tris-HCl pH 7.5, 10 mM magnesium chloride, 20% dimethylformamide). After washes, beads were resuspended in 25 µL of 2x TD Buffer, Tn5 (for 50 ng of post-ChIP DNA we used 2.5 µL of Tn5), and water to 50 µL. Tn5 amount was adjusted linearly for different amounts of post-ChIP DNA, with a maximum amount of 4 µL of Tn5. Samples were incubated at 55°C with interval shaking for 10 min. After removing the supernatant 50 mM EDTA was added to samples and incubated with interval shaking at 50°C for 30 min. Beads were then washed two times each in 50 mM EDTA then Tween Wash Buffer at 55°C for 2 min. Lastly, beads were washed in 10 mM Tris before PCR amplification. Beads were resuspended in 25 µL of Phusion HF 2x (New England Biosciences), 1 µL of each Nextera Ad1_noMX and Nextera Ad2.X at 12.5 µM, and 23 µL of water. The following PCR program was performed: 72°C for 5 min, 98°C for 1 min, then cycle at 98°C for 15 s, 63°C for 30 s, and 72°C for 1 min (cycle number was estimated based on the amount of material from the post-ChIP Qubit (approximately 50 ng was run in six cycles, while 25 ng was run in seven, 12.5 ng was run in eight, etc.). Size selection was performed using two-sided size selection with the Ampure XP beads. After PCR, libraries were placed on a magnet and eluted into new tubes. 25 µL of Ampure XP beads were added, and the supernatant was kept to capture fragments less than 700 bp. Supernatant was transferred to a new tube, and 15 µL of fresh beads was added to capture fragments greater than 300 bp. After size selection, libraries were quantified with Qbit and sent for Bioanalyzer to check for the quality and final size of the library. Libraries were sequenced on a HiSeq 2500 platform on PE75 mode.

### 4C-seq

For each sample 10 million cells were fixed following our ChIP-seq protocol (see above). Cell pellets were lysed in 1ml Lysis Buffer (50 mM Tris-HCl pH 7.5, 150 mM NaCl, 5 mM EDTA; 1x complete protease inhibitor, 0.5% NP-40, 1% triton) and incubated on ice for 15 min. The samples were centrifuged at 2500xG for 5min at 4°C and the pellet was then resuspended in 360µl milli-Q, 60µl 10X DpnII restriction buffer and 15ul 10%SDS. After 1 hour incubation at 37°C, 150ul of 10% Triton was added and samples were incubated again at 37°C for 1 hour. 4ul DpnII enzyme (#R0543M, NEB) were added and samples were incubated at 37°C over night while shaking in a thermomixer (9000rpm). After confirming the digestion efficiency, the enzyme was inactivated by adding 80ul 10% SDS and incubating at 65 °C for 30 mins. The digested samples were then diluted with 4860ul Milli-Q, 700ul ligation buffer (500mM Tris pH 7.5, 100mM DTT, 100mM MgCl2,10mM ATP), and 750ul of Triton and incubated at 37°C for 1 hour. Then 2ul Ligase (NEB M0202M) were added and samples were incubated over night at 16 °C. Next morning, after testing the ligation efficiency, we reversed the crosslinks by adding 30ul of proteinase K (10mg/ml) and incubating over night at 65°C. Subsequently the RNA was removed using 30ul of RNase A (10mg/ml) for 45mins at 37°C. Extensive phenol/chloroform extraction was followed by EtOH precipitation and two washes with 70% EtOH. The pellets were dissolved in 150ul 10mM Tris pH 7.5 by incubating at 37 °C. We then added 50ul 10x buffer B (Fermentas), 5ul Csp6I (Fermentas, ER0211) and 299ul milli-Q water and samples were digested at 37°C over night. After determining digestion efficiency, the restriction enzyme was inactivated by incubating the tubes at 65°C for 25 mins. Samples were diluted in 12ml milli-Q, 3ul ligase (NEB, M0202M) and 1.4ml 10X ligation buffer (500mM Tris pH7.5, 100mM DTT, 100mM MgCl2, 10mMATP) and incubated over night at 65°C. Following phenol/chloroform and EtOH precipitation the pellets were dissolved in 300ul 10mM Tris pH7.5 and DNA was further purified using 4 Zymo columns per sample (Zymo, D4014). Each sample was eluted in 200ul total of 10mM Tris pH7.5. Finally, 150ng of DNA was used per reaction, to PCR-amplify the libraries using the KAPA HiFi enzyme (KAPA biosystem, 07958927001). All primer sequences can be found in Supplementary table 9. Four PCR reactions were combined per sample, following column purification using the ZYMO kit (Zymo, D4014). Samples were sent for QC on a bioanalyzer and then sequenced on a HiSeq 4000 platform on SE50 mode.

### RIME

RIME was performed in 3 replicates for KLF4 and 2 for IgG, as previously described^52^ with minor modifications. 50 million V6.5 cells grown in 2i conditions were used for each replicate. Cells were fixed, lysed, sonicated and incubated with the respective antibody-bound beads, using the same conditions that were used for KLF4 ChIP-seq (see above). The samples were then washed ten times in RIPA buffer (50 mM HEPES (pH 7.6), 1 mM EDTA, 0.7% (wt/vol) sodium deoxycholate, 1% (vol/vol) NP-40 and 0.5M LiC) and five times in 100mM AMBIC solution. Treatment for enzymatic digestion and peptide desalting was carried out as in the original protocol.

### Co-IP and WB

50 million V6.5 cells grown in 2i condition were collected for each Co-IP experiment and resuspended in 0.5ml lysis buffer (50mM Tris pH7.5, 100mM Nacl, 0.2% triton, 0.5% glycerol and protease inhibitors). Cells were incubated on ice for 40 mins followed by 3 cycles of sonication in a bioruptor device (30sec on/off, high setting) and spun down 10 minutes at 4°C at maximum speed. Supernatants were diluted with additional lysis buffer in a final volume of 2ml. Lysates were pre-cleared with 10ul of protein G Dynabeads (ThermoScientific) for 30 mins in rotation at 4°C. The supernatant was then incubated with 8ug of KLF4 antibody (R&D, AF3158) or IgG (Calbiochem, NI02) for 2.5 hours in rotation at 4°C. 30ul of protein G Dynabeads that were pre-blocked with BSA were added to the samples and incubated 1.5hours in rotation at 4°C. Two washes were performed with lysis buffer followed by three washes with high salt buffer (same as lysis buffer but with 250mM NaCl). Finally, the samples were eluted in loading buffer by boiling 5 minutes and transferring the sup to a new tube. WBs were performed with the following antibodies: BRD4 (Bethyl, A301-985A50), MED1 (Bethyl, A300-793A), SMC1a (Bethyl, A300-055A), RING1b (Bethyl, A302-869A), SUZ12 (Santa Cruz, sc46264), LSD1(Abcam, ab 17721).

### ATAC-seq data analysis

*Mapping, peak calling and peak processing*. Paired-end reads were aligned to mm10 (bowtie2 version 2.3.2, --no-unal --local --very-sensitive-local --no-discordant --no-mixed --contain --overlap --dovetail -I 10 -X 2000), and mitochondrial DNA alignments were excluded. Fragments marked as positional duplicates (sambamba version 0.6.6) or overlapping with mouseENCODE blacklisted genomic regions^80^ (liftOver to mm10) were filtered out. Read ends were adjusted for Tn5 transposase offsets. Peaks were called at p<10^-5^ (MACS version 2.1.1) per replicate, and only common peaks between two independent biological replicates were retained for further analysis.

### ChIP-seq data analysis

*Mapping, peak calling and peak processing*. Study and published ChIP-seq reads were trimmed for adapters (cutadapt version 1.8.1), and low-quality ends (sickle version 1.33), respectively. Alignment to the mouse reference genome version mm10 (GRCm38.p4) was performed using standard parameters, permitting a maximum of one mismatch in seed alignment (bowtie2 version 2.3.2). Reads marked as positional duplicates (sambamba version 0.6.6) or overlapping with mouseENCODE blacklisted genomic regions (liftOver to mm10) were filtered out. Study ChIP-seq peaks (enrichment of signals over background determined by input samples) were called at p<0.01 (MACS version 2.1.1) per biological replicate, and peaks detected in more than half of biological replicates were retained for further analysis. Published ChIP-seq replicates were merged, and peaks were called at p<10 ^-5^ using input samples where applicable.

### Overlap analysis of ChIP-seq peaks for chromatin states of reprogramming cell types

Chromatin states (1 kb resolution) during reprogramming were retrieved from ref^8^, and cis-regulatory elements were annotated from chromatin states as in the original publication. The assignment of ChIP-seq peaks to cis-regulatory elements was determined by the largest degree of overlap in bp.

### RNA-seq and ChIP-seq gene ontology (GO) enrichment analysis

Spatial proximity of ChIP-seq peaks to transcript start sites (TSSs) and enriched GOs were uncovered utilizing the GREAT (version 3.0.0) web application. We selected the “basal plus extension rule” for the association of gene ontology annotations with regulatory domains (customized setting: 5 kb upstream and 1 kb downstream of TSSs, and further extended both directions by 250 kb). Enrichment of ontology annotations was assessed by the binomial test of ChIP-seq peak-overlaps with annotated regulatory regions. For differentially expressed genes and gene groups (Fig. S4e) DAVID knowledgebase^81^ was used for pathway and biological process enrichment analysis.

### PSC typical enhancers and super-enhancers

Coordinates of typical- and super-enhancers in mESCs and other cell lines or tissues were ascertained from ref^45^ and ref^34^, lifted over from mm9 to mm10 with UCSC liftOver.

### Overlaps of KLF4 binding with early lost or late gained H3K27ac peaks during reprogramming or at typical- and super-enhancers

The maximum permitted distance between KLF4 binding detected in day3, 6 and 9 with PSC and H3K27ac peaks or enhancers in ref^45^ was 250 bp. Where H3K27ac peaks or enhancers overlapped with KLF4 sites of different stages, the earliest stage was prioritized (Fig. 1g).

### Motif analysis

For each KLF4 cluster we generated a random background (by shuffling the peaks randomly throughout the genome) to test motif enrichment within each cluster. Analysis of the KLF4 clusters was performed with the use of HOMER and ‘findMotifsGenome.pl’ command with the following parameters: ‘-bg random.bed - size 200 -len 15’. Only motifs with p-value≤1e-5 were considered significant. Two heatmaps with the z-transformed ‘-log10(p-value)’ and z-transformed ‘motif frequency’ of selected motifs for each cluster are presented in Supplementary Fig.1e.

### PCA analysis for ATAC-seq, RNA-seq and ChIP-seq experiments

We first merged all accessible regions / H3K27ac peak detected from ATAC-seq / H3K27ac ChIP-seq in any reprogramming stage using bedtools v2.25.0. Then, we calculated the coverage of reads for each merged accessible region and H3K27ac peak for each replicate independently. For the RNA-seq data, we calculated the coverage for each exon and only exons with at least 1 read covering every single base of the exon were used for downstream analysis. PCA analysis was performed with R and PCA plots were generated with ‘ggplot2’ library. In each PCA plot, we present the variability captured by the first two PCs (PC1 and PC2).

### RNA-seq data analysis

Expression of genes was quantified in transcripts per kilobase million (TPM) using quasi mapping (Salmon, version 0.8.2) to GENCODE (version M6, mm10) reference gene annotation. Salmon provides alignment-free transcript quantification information in a single step^82^.

### Line plots for gene expression analysis

We plotted the median expression levels of all protein coding genes with their corresponding 95% confidence interval (CI) that are bound by KLF4 in a distance less than 50 kb from their corresponding transcription start site (TSS). For each KLF4 cluster we calculated the closest (≤50 kb) TSS to each KLF4 binding site and plotted the median expression levels (TPM) of all genes annotated in each KLF4 cluster with the use of R.

### Processing of HiChIP / HiC datasets

HiChIP and HiC datasets were uniformly pre-processed with the HiC-bench platform^83^, which is outlined in short in the following. First, all paired-end sequencing reads were aligned against the mouse genome version mm10 with bowtie2 version 2.2.3^84^ (specific settings: --very-sensitive-local --local). Read-filtering was conducted by the GenomicTools^85^ gtools-hic filter command (integrated in HiC-bench), which discards multi-mapped reads (“multihit”), read-pairs with only one mappable read (“single sided”), duplicated read-pairs (“ds.duplicate”), read-pairs with a low mapping quality of MAPQ < 20, read-pairs resulting from self-ligated fragments and short-range interactions resulting from read-pairs aligning within 10kb (together called “ds.filtered”). The percentage of accepted intra-chromosomal read-pairs (“ds.accecpted intra”) was high across all HiC and HiChIP replicates and conditions and was consistently above 35%. In order to create counts-matrices per chromosome in a binned fashion, we set the bin size to 10kb for all datasets. For all the HiChIP sample and chromosome matrices, the trajectories of each matrix bin to both anchors were overlaid with the ChIP-Seq signal of the respective matching sample, requiring a minimal overlap of 1bp between a HiChIP-bin and a ChIP-peak. Only loops of which at least one anchor was supported by a ChIP-peak were kept for further analyses. Next, we applied sequencing-depth normalization (leading to read-counts per million, or CPM) per replicate followed by a statistical approach to identify significant loops. We have adapted the approach first described in Mango^37^, by performing a binomial test in each diagonal of the counts-matrix up to a maximum distance of 2MB.

High-confidence HiChIP loops were identified by p-value < 0.1 and requiring a CPM > 3 per loop across all replicates of a single condition in order to maintain a signal that is replicable. For high-confidence HiC loops, we have adjusted those thresholds in order to avoid too much noise, and have applied filters of p-value < 0.01 and CPM > 15 across all replicates of a single condition.

### Principal component analysis for HiChIP samples

Principal component analysis (PCA) as shown in Figures S5a was performed on all available replicates on the high-confidence loops. Therefore, for each detected high-confidence loop from any sample, the per replicate normalized CPM was extracted before filtering for significant loops in order to also integrate lowly detected interactions in the analysis. PCA was performed using the prcomp function of R (version 3.3.0; scale=TRUE, center=TRUE).

### Differential loop analysis

Differential looping analysis was performed on each significant loop independently by applying an unpaired two-sided t-test on the normalized counts (CPM) calculated before identifying significant loops between any pairwise comparisons: PSC-KLF4 vs d3-KLF4, PSC-KLF4 vs d6-KLF4, d3-KLF4 vs d6-KLF4, PSC-H3K27ac vs MEF-H3K27ac, TKO-0h vs TKO-24h. In order to estimate the change in loop strength, we calculated the log2 fold-change (logFC) between the average CPM per condition for the same pairwise comparisons after adding a pseudo-count of 1 to each replicate and loop. For constant H3K27ac loops in either MEF vs PSC or TKO-0h vs TKO-24h, we selected loops with p-value > 0.5 and absolute logFC < 0.5 for the respective pairwise comparison. MEF/PSC-specific H3K27ac loops were selected by p-value < 0.1 and logFC > 2 / logFC < −2 taken from the PSC H3K27ac vs MEF H3K27ac comparison, respectively. TKO-0h/TKO-24h specific loops were selected by p-value < 0.05 and logFC > 0.58 / logFC < −0.58 taken from the TKO-0h vs TKO-24h comparison. Mid and late established KLF4 loops were selected by applying p-value < 0.01 and logFC > 2 in the pairwise comparisons of PSC-KLF4 vs d3-KLF4 and d6-KLF4 vs d3-KLF4 (mid) and PSC-KLF4 vs d3-KLF4 and PSC-KLF4 vs d6-KLF4 (late). Early-lost and mid-lost KLF4 loops were selected by applying a p-value < 0.01 and logFC < −2 in the pairwise comparisons of PSC-KLF4 vs d6-KLF4 and PSC-KLF4 vs d3-KLF4 (early-lost) and PSC-KLF4 vs d3-KLF4 and d6-KLF4 vs d3-KLF4 (mid-lost). For differential comparison of significant HiC loops, we have applied a distance-normalization as previously described^86^ before calculating significance and fold-changes between PSC and MEF HiC loops. Then, differential HiC loops were selected by applying a p-value < 0.1 and logFC < −0.32 or logFC > 0.32 (equivalent to a fold-change of 1.25) in the pairwise comparison of PSC-HiC vs MEF-HiC. All calculations were performed in R version 3.3.0, using the native t.test function (unpaired, two-sided).

### Annotation of H3K27ac HiChIP loop anchors as promoters or enhancers

H3K27ac HiChIP loop anchors were overlapped with transcription start sites (TSSs) of GENCODE (version M6) protein coding genes. Presence of one or more TSSs was considered a promoter HiChIP anchor, and the absence of any TSS but presence of at least one H3K27ac ChIP-seq constitutes an enhancer HiChIP anchor. In estimating connectivity, all HiChIP anchors, either promoter, enhancer or otherwise desolate, were considered.

### RNA expression integration with differential HiChIP loops

For RNA expression integration, we overlapped all canonical TSSs of protein-coding genes (transcript support level/TSL = 1) downloaded from Ensembl Genes V85 for the mouse genome mm10 with all loop anchors. Because the TSS is a 1bp position in the genome, each gene was uniquely assigned to one bin, however, multiple TSSs per gene with a TSL=1 mapping to different bins are possible. Next, we filtered genes by occurrence of differential loop clusters that were obtained from the HiChIP experiments and have TPM > 1 in at least one reprogramming stage, and analyzed the expression patterns of such genes throughout reprogramming. For H3K27ac HiChIP data integration, we assigned genes to MEF/PSC-specific loops if their TSSs were found in >= 1 MEF/PSC-specific loops but in none of the other (Figure 2b). Genes contained in anchors of constant loops were filtered by having at least 1 or 3 constant loop anchors but no MEF/PSC-specific loop. To further validate expression changes based on differential looping, we applied an unpaired, one-sided t-test between genes logFCs of constant H3K27ac loops versus genes with MEF/PSC-specific loops, following the hypothesis of a positive correlation between looping and expression changes. As a negative control, we compared logFCs of genes with constant loops versus all annotated protein-coding genes. We have followed the same approach for the integration of expression data with differential loops obtained from Klf-TKO H3K27ac HiChIP experiments. In short, we assigned genes to TKO-0h/TKO-24h specific loops if their TSSs were found in >= 1 differential loops but not in the other differential loop category. We have compared logFCs of expression of TKO-0h and TKO-24h between TKO-0h/TKO-24h specific and constant loops.

### Co-regulation of gene expression by H3K27ac HiChIP (enhancer hubs)

In this analysis, promoter anchors of enhancer-mediated loops were filtered for protein-coding genes that have an expression TPM > 1 in PSC. Enhancer fragments that contact two to ten promoter fragments in PSC specific H3K27ac loops were selected. Genes were paired across different promoter fragments connecting to the same enhancer anchor (later on called hub), and repeated gene pairs were removed from the overall pool. Gene pairs were considered co-expressed, if both genes were up-regulated in PSC compared to MEF (p-adjusted<10^-2^ and fold change threshold of 2) or vice versa for down-regulation. Or otherwise, at least one gene in a pair unchanged between MEF and PSC constitute unchanged gene pairs. In order to test if the enrichment of the co-regulated gene pairs in the original hubs was significant we performed Fisher’s exact test. The background probability was calculating by using an equal number of random gene pairs (protein-coding genes that have an expression TPM > 1 in PSC) either of similar linear distance with our test group (global random) or within the same TADs. TADs were called from normalized corrected HiC matrices in PSCs processed at 10kb resolution using a recently published software^87^ with the use of the following parameters ‘--minDepth 120000 --maxDepth 420000 -- thresholdComparison 0.001 --delta 0.01 --correctForMultipleTesting fdr’.

### Overlap between H3K27ac loop clusters and KLF4 clusters

Overlap between any of the KLF4 peaks with any of the HiChIP anchors (H3K27aC or KLF4 loops) was performed with the use of bedtools v2.25.0. Odds ratio and significance of the overlap between the 2 groups was performed with the use of Fisher’s exact test.

### ChIP-seq feature enrichment at lowly or highly connected H3K27ac PSCs specific enhancer anchors

H3K27ac HiChIP enhancer anchors were selected for low (N = 1,183) or high connectivity (contacting four or more anchors; N = 1,014). LOLA analysis was performed in these two groups of ChIP-seq peaks in order to identify which TFs participate in the formation of low vs high connected hubs (Fig.3g).

### KLF4 looping involved in RNA expression changes

To estimate the effect of KLF4 associated looping on changes in RNA expression, we followed a similar approach as before for the H3K27ac HiChIP integration. After selecting expressed genes within anchors of each KLF4 loop cluster, we further filtered for differentially expressed genes between PSC and day3 (FDR < 0.01; logFC > 1.0 (upregulated) or logFC < −1.0 (downregulated)). Information on differential expression was derived by DESeq with subsequent multiple testing correction as mentioned before. Genes determined as ‘no change’ were selected by applying FDR > 0.5 and absolute logFC < 0.25. All remaining genes were discarded from the analysis.

### LOLA enrichment analysis

The identified differential loops were subjected for an enrichment analysis of further transcription factor bindings and other DNA binding proteins. First, the anchors of each differential loop were mapped back to the original ChIP or ATAC-peaks, because the 10kb stretches of the bins would give too many false positive findings. Each anchor that was overlapping an actual ChIP or ATAC-peak by at least 1bp was subjected for further analysis. Since two anchors can theoretically overlap with a single ChIP-peak using this approach, the resulting list was collapsed and only unique ChIP or ATAC-peaks were kept. Next, we applied LOLA version 1.8.0^88^ against a database of analyzed ChIP-Seq datasets taken from LOLA Region Databases (regionDB) for mm10 (for Figure S3e we used Codex and encode TFBSmm10 databases). We excluded all ChIP-Seq datasets that were marked as treated with any agent and had less than 3000 peaks in total. When multiple ChIP-seq data for the same antibody were significantly enriched in one of our tested regions we selected the one with the highest number of peaks. In addition to the ChIP-seq peaks provided by the LOLA database we manually constructed a database containing ChIP-seq from the following studies GSE22557, GSE90893, GSE99519, GSE22562 and our own ChIP-seq data. Data from these studies were re-analyzed with the same pipelines that were used for our ChIP-seq data. As a universe for LOLA, we used only unique ChIP or ATAC-peaks from the union of all ChIP or ATAC-Seq peaks for the respective antibody across all reprogramming stages.

### Virtual 4C

Virtual 4C was performed to identify interaction signals of gene promoters or enhancers with their genomic vicinity. For this approach, we used the filtered HiChIP read-pairs as described above before binning and normalization of each replicate. We extracted all read pairs for which a read mate maps within +/-10kb around the virtual viewpoint. Next, we defined successive overlapping windows for each chromosome at a 10kb resolution, and all adjacent windows are overlapping by 95% of their length (i.e. 9.5kb, or a shift of 500bp between adjacent windows). We then counted the second mapped read mate in all overlapping bins. Thus, each read-pair accounts for +1 in exactly 19 bins, however, the overlap of bins achieves a smoothed signal. Read counts for all bins were normalized to total sequencing depth of the respective replicate by edgeR version 3.14.0 to calculate counts-per-million (CPM) per bin. Significant differences between any condition (TKO-0h vs TKO-24h H3K27ac HiChIP or MEF vs ES H3K27ac HiChIP) was calculated using edgeR function glmQLFTest (we have not corrected for multiple testing, because the requirement of independent data-points for multiple testing correction is not given for the overlapping windows). For visualization, the average of the normalized virtual 4C-signal across replicates of a single condition was calculated.

### Analysis of 4C-seq data

The 4C-seq data was analyzed in a similar fashion as recently described^89, 90^. Firstly, viewpoint primers were trimmed off from all sequencing reads using seqtk (version 1.3.0). Next, the remaining read-sequence was aligned using bowtie v1.0.0 against a reduced genome that consists only of reference genome sequences adjacent to DpnII cut-sites which was used during the 4C protocol (following the 4C-ker pipeline^89^). By aligning against the reduced genome, only reads matching the adjacent sequence of an actual digestion fragment are allowed, and the remaining reads are automatically discarded. Next, the genome was binned into 10kb bins shifted by 500bp (thus overlapping by 95% with adjacent bins), similar as the virtual 4C approach described above. Reads were counted by unique alignment position per bin, thus accounting for +1 read in 19 adjacent bins to achieve a smoothed signal. Read counts per bin were normalized by sequencing depth per replicate using edgeR (version 3.14.0), resulting in counts per million (CPM). The visualization shows the average CPM signal across all replicates of a single condition.

### RIME analysis

Summed ‘signal to noise’ intensity per protein from 3 KLF4 and 2 IgG samples was used to calculate significant enrichment of KLF4 protein complexes with the use of Welch’s t-test. Only proteins with a p-value <0.05 and fold enrichment greater than 1.5 over IgG were considered significantly enriched in our samples.

---

**Supplementary Figure 1.**
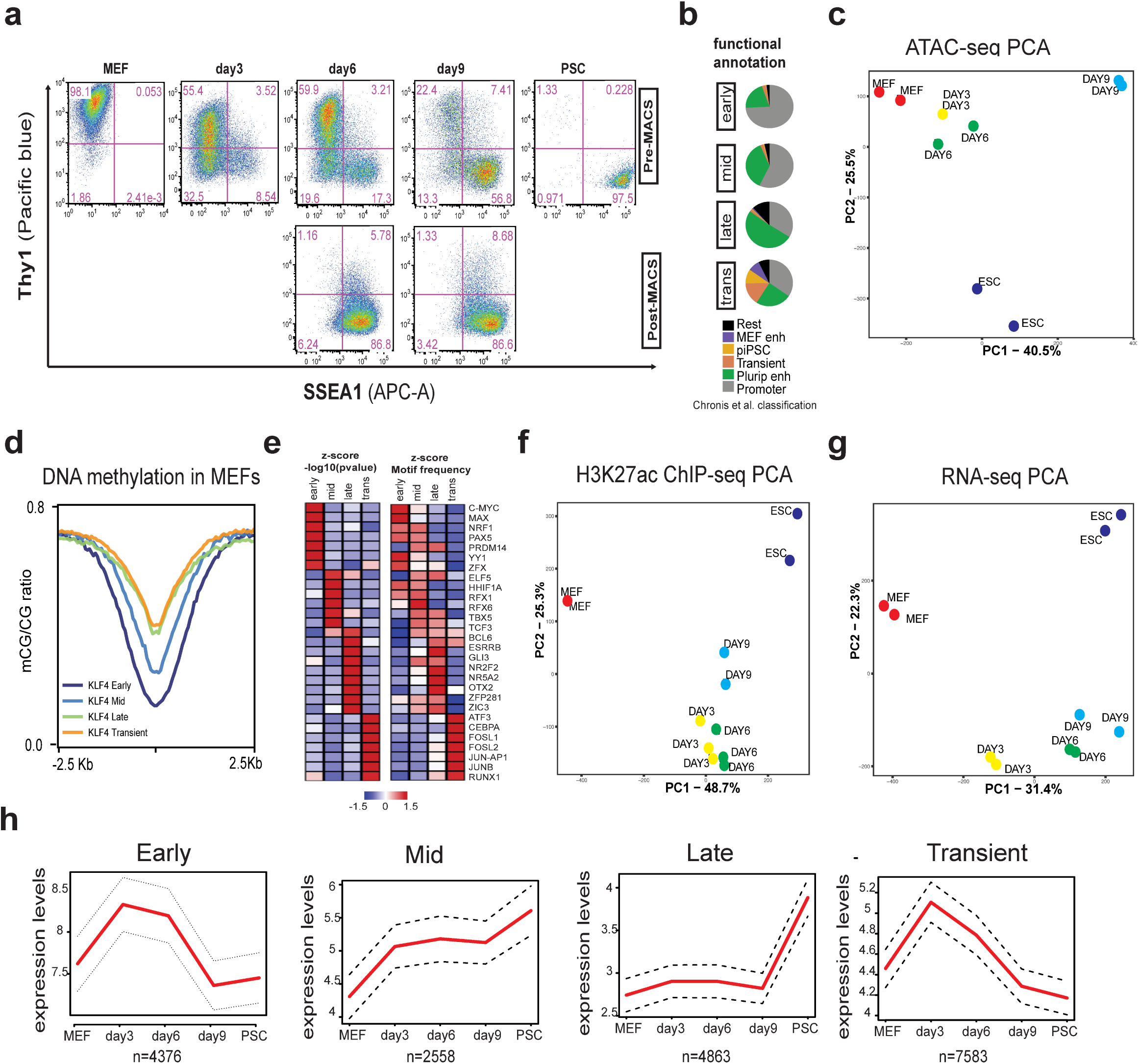
**a**, FACS analysis plots showing expression of SSEA1 (early pluripotency marker) and Thy1 (somatic marker) at different stages of reprogramming, before and after SSEA1 enrichment by MACS isolation. **b**, Pie charts of functional classification of KLF4 Early, Mid, Late and Transient peaks (based on Chronis et al. 2017) (piPSC= partial iPSCs). **c**, PCA analysis of ATAC-seq peaks in MEF, PSC and different stages of reprogramming. **d**, Average line plot showing the methylated CG to non-methylated CG ratio from MEF data^12^ centered (+/-2.5Kb) around different clusters of KLF4 binding sites (Early, Mid, Late or Transient KLF4 targets, Fig.2b). **e**, Motif enrichment for Early, Mid, Late and Transient KLF4 binding sites. Selected factors are shown and their significance is expressed as Z-score of –log10(pvalue) (left) or z-score of motif frequency (right). **f**, PCA analysis of H3K27ac ChIP-seq peaks called in MEF, PSC and different stages of reprogramming **g**, PCA of RNA-seq in MEF, PSC and different stages of reprogramming. **h**, Line plots of the median expression (red line) of genes closest to Early, Mid, Late and Transient peaks, expressed as TPM (transcripts per million).

**Supplementary Figure 2.**
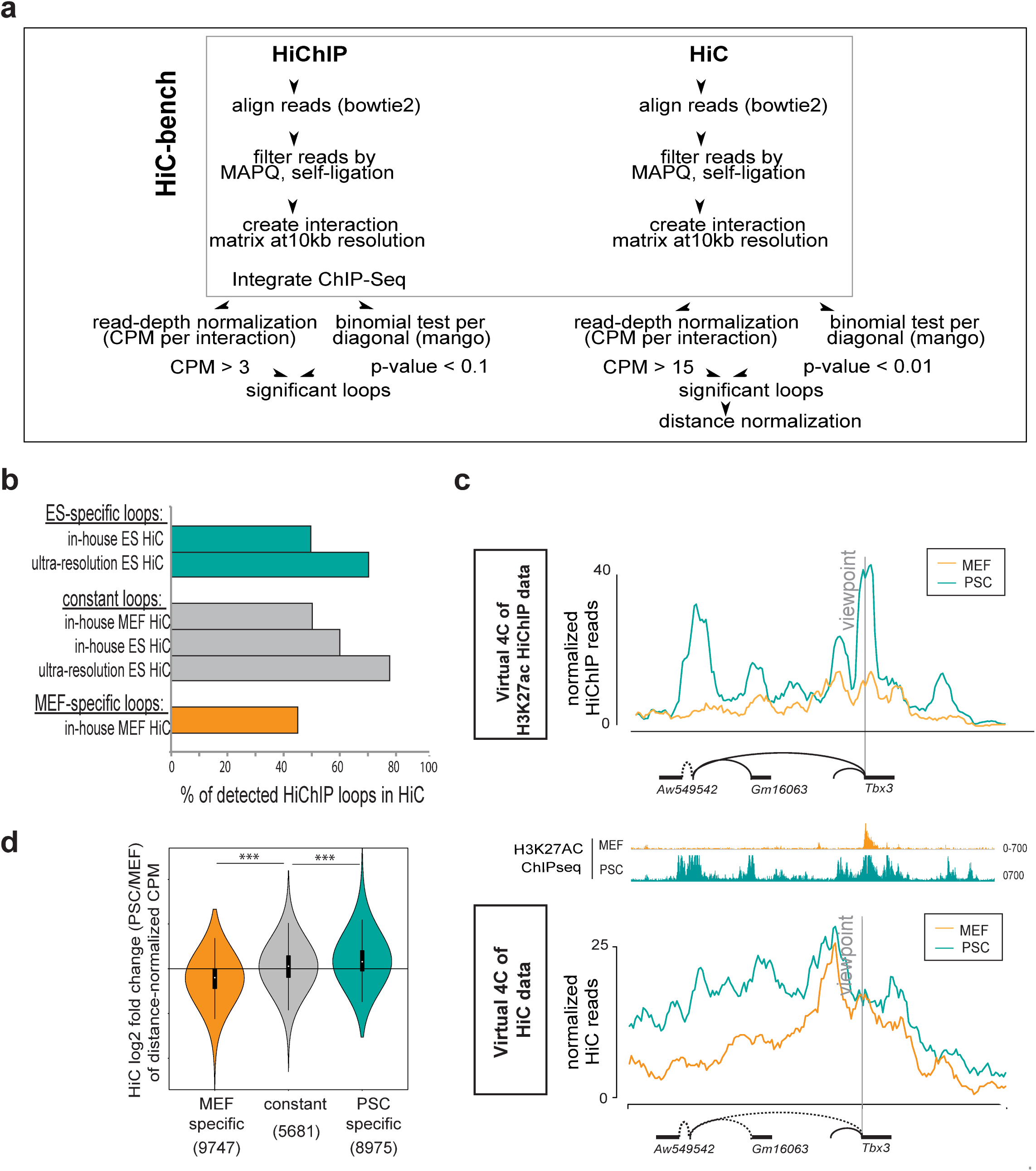
**a**, Schematic work-flow for HiChIP and HiC analysis. **b**, Percentages of PSC-specific, constant or MEF-specific H3K27ac HiChIP loops that were detected in HiC experiments (either generated in-house or published ultra-resolution HiC in PSC^38^). **c**, Normalized HiChiP (top) and HiC (bottom) signals in MEF and PSC are illustrated in a virtual 4C format around the indicated viewpoint (*Tbx3* promoter). H3K27ac ChIP-seq tracks are shown in MEF and PSC. **d**, Violin plot representing log2 fold change of distance-normalized HiC signal in PSCs versus MEFs of MEF-specific, constant and PSC-specific loops as called by H3K27ac HiChIP. Only contacts that were detected as significant in HiC data are considered. Numbers of considered loops per category are shown in parenthesis. Unpaired two-sided t-test was used to determine the p-value.

**Supplementary Figure 3.**
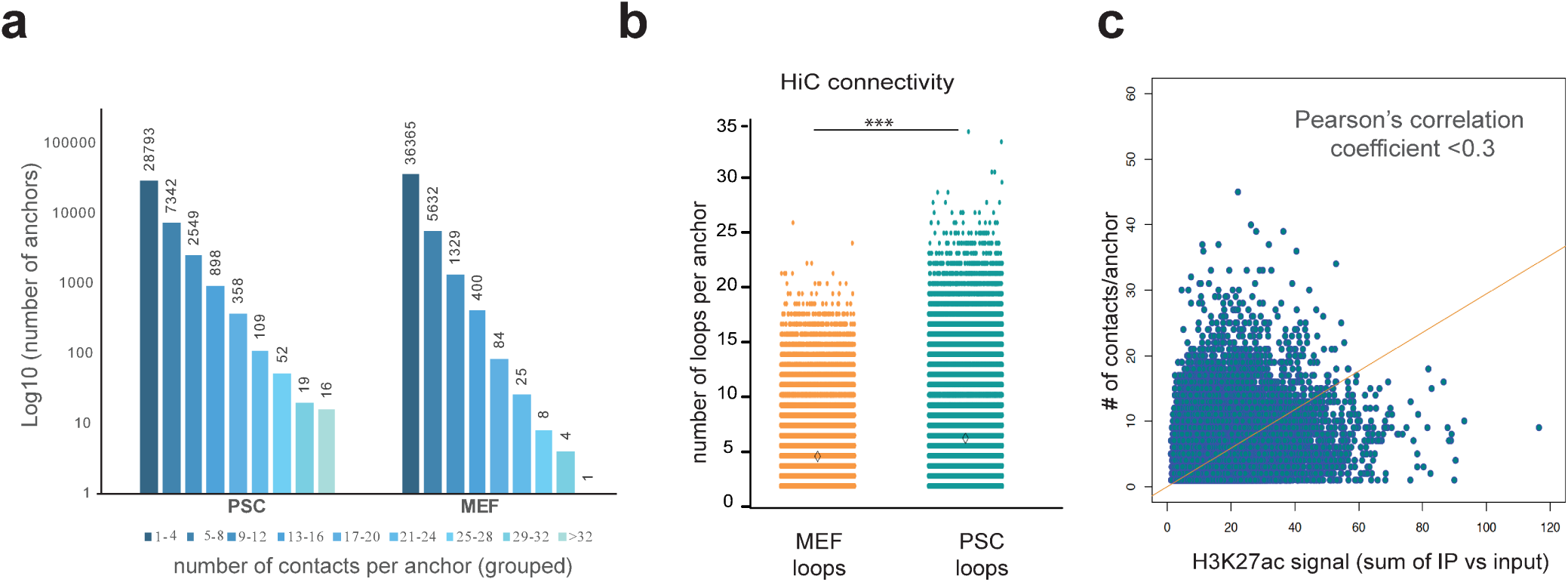
**a**, Histogram of anchor connectivity based on H3K27ac MEF and PSC HiChIP called loops. The numbers of contacts per anchor are grouped as shown in the bottom and the actual number of anchors is depicted on top of each bar. **b**, Connectivity of MEF or PSC anchors based on HiC-called loops represented as number of high-confidence contacts around each 10kb anchor. Wilcoxon rank sum test was used to compare connectivity and asterisks indicate significant difference with p<0.001. **c**, Scatter plot showing the correlation of H3K27ac ChIP-seq strength (sum of H3K27ac ChIP/input of all peaks within the anchor) with the number of H3K27ac HiChIP contacts per anchor in PSCs.

**Supplementary Figure 4.**
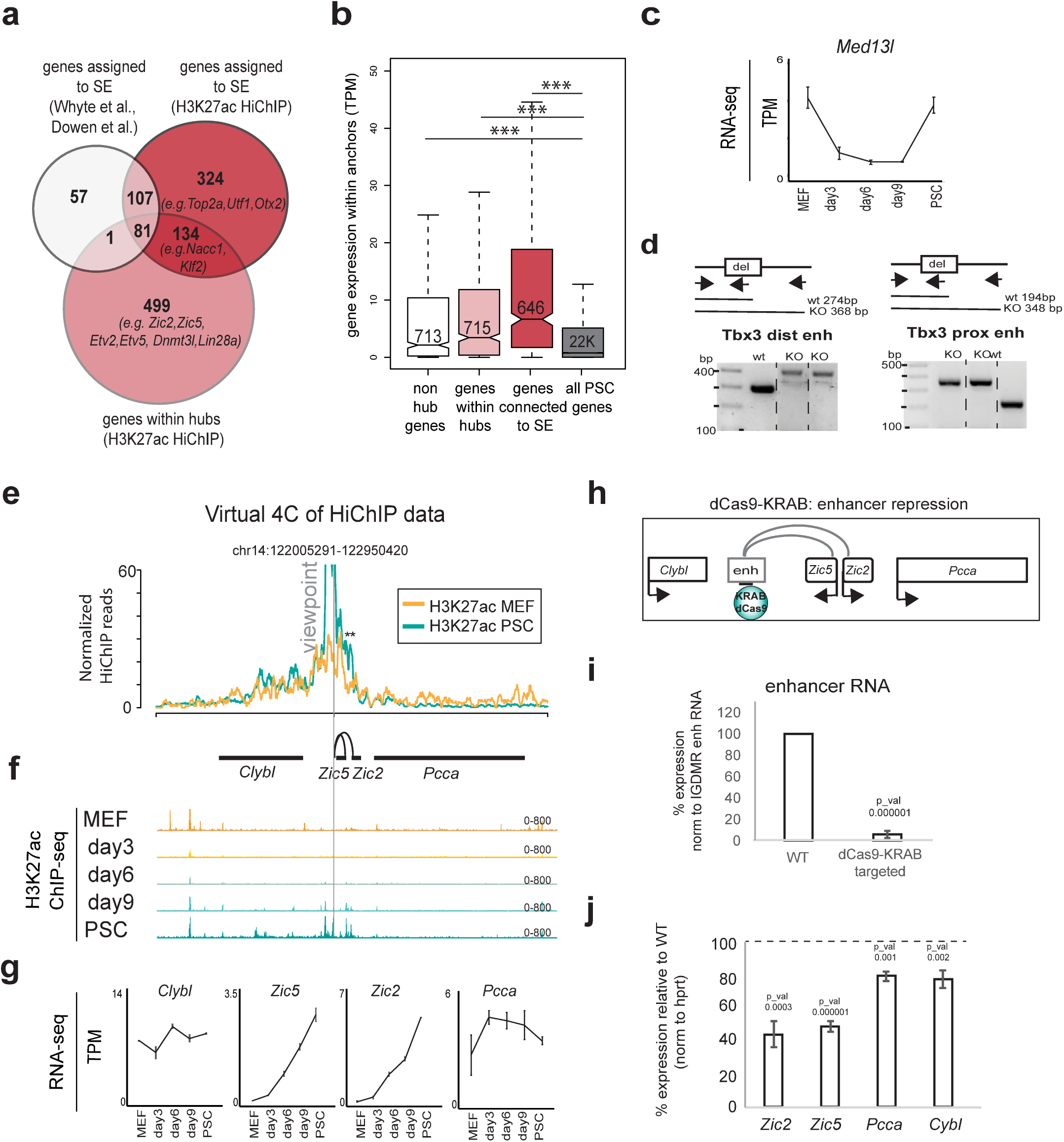
**a**, Venn diagram showing overlap between previously assigned target genes for super-enhancers (SE), newly identified SE target genes based on H3K27ac HiChIP contacts in PSCs, and genes connected to PSC-specific enhancer hubs, which represent enhancers contacting more than one gene according H3K27ac HiChIP (see also Fig.4a). **b**, RNA levels of hub genes, non-hub genes or genes connected to SE in PSC samples as measured by RNA-seq and expressed as transcripts per million (TPM). All genes that are not connected to enhancer hubs, but are still detected within PSC-specific HiChIP loops were considered. Expression of all genes expressed in PSC (>1TPM) is shown as reference. **c**, RNA-seq signal (TPM) of *Med13l -*which is not part of the *Tbx3* enhancer hub (see Fig.4b)- during reprogramming **d**, Genotyping strategy and results confirming the homozygous deletion of the distal (left) or the proximal (right) *Tbx3* enhancers. **e**, Example of a newly identified enhancer hub in PSCs. Normalized HiChIP signal around the viewpoint is illustrated as a virtual 4C plot. **f**, H3K27ac ChIP-seq IGV tracks during reprogramming. **g**, RNA-seq signal of genes within the hub (*Zic2* and *Zic5*), or nearby genes (*Clybl* and *Pcca*), are shown for each reprogramming stage to highlight concordance with H3K27ac HiChIP data and coordinated upregulation of genes within the hub. **h**, Schematic illustration of the CRSIPRi (dCas9-KRAB) targeting strategy for inactivation of the *Zic2/Zic5* enhancer hub. **i**, RT**-**qPCR showing relative levels of the enhancer RNA (normalized to an unaffected enhancer RNA (IGDMR)) in wild type (WT) or dCas9-KRAB-targeted ESCs. P-values were calculated using unpaired one-tailed t-test. Error bars indicate standard deviation from n=2 biological replicates. **j**, RT**-**qPCR showing expression changes of genes within the hub (*Zic2* and *Zic5*) and nearby genes (*Clybl* and *Pcca),* calculated as percentage relative to WT after normalization to *Hprt* expression. P-values were calculated using unpaired one-tailed t-test. Error bars indicate standard deviation from n=2 biological replicates.

**Supplementary Figure 5.**
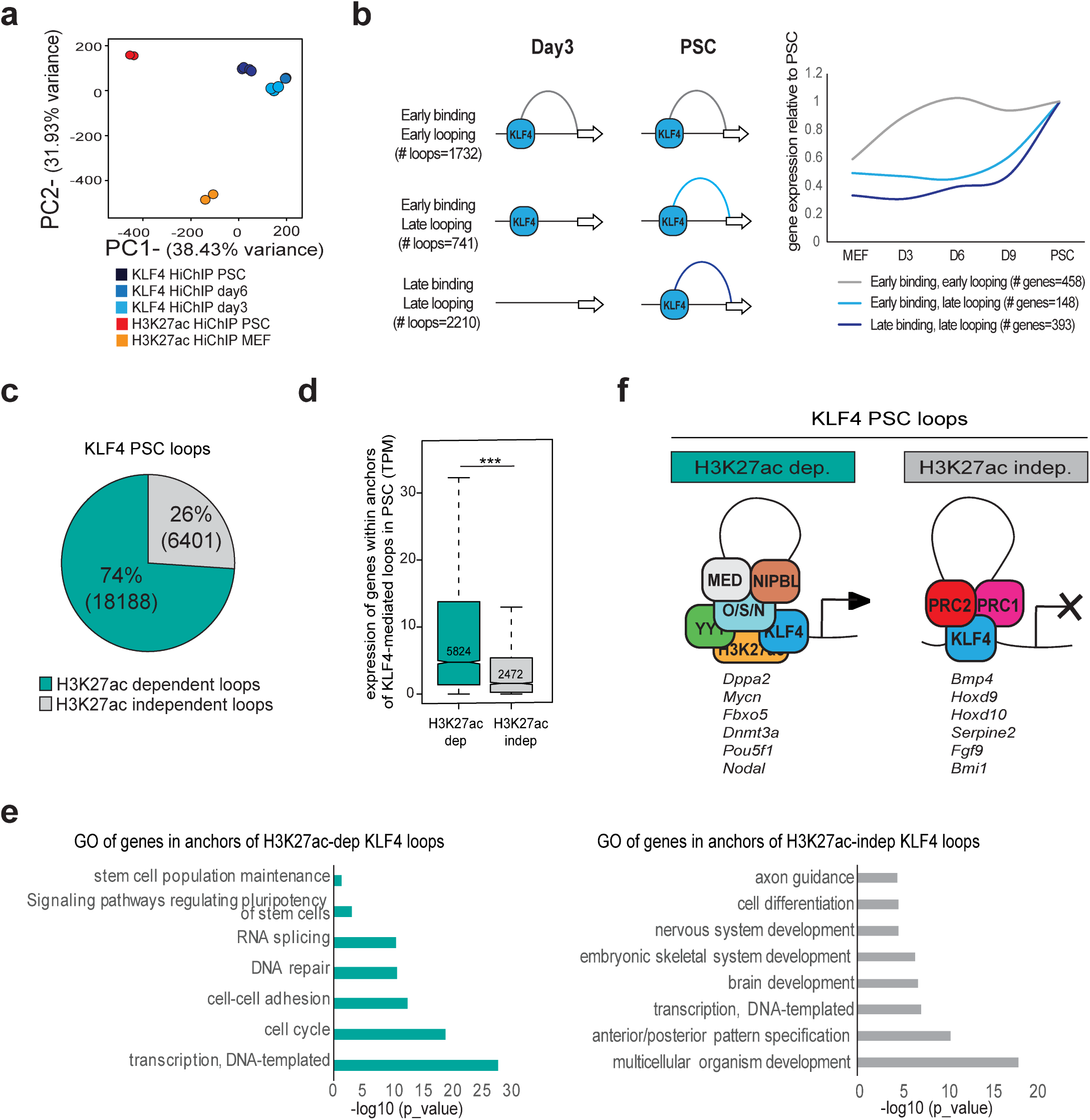
**a**, PCA analysis of loops called as significant by H3K27ac and KLF4 HiChIP in different samples. **b**, Left: Chromatin loops that were detected by both KLF4 and H3k27ac HiChIP in PSCs were clustered based on the timing of KLF4 binding and looping during reprogramming. Right: Line plot showing expression changes of genes that belong to each of the indicated loop categories during reprogramming (median values are plotted relative to PSC). **c**, Pie chart showing the percentage of KLF4 PSC loops that were also detected by H3K27ac HiChIP in PSCs (H3K27ac-dependent) or not (H3K27ac-independent). **d**, Boxplot showing expression of genes within all anchors of KLF4-mediated loops that are either H3K27ac-dependent or independent. **e**, Gene ontology for genes within anchors of H3K27ac-dependent or -independent KLF4 loops. **f**, Proposed model for different categories of chromatin loops mediated by KLF4 and cofactors. Example genes are reported for each category.

**Supplementary Figure 6.**
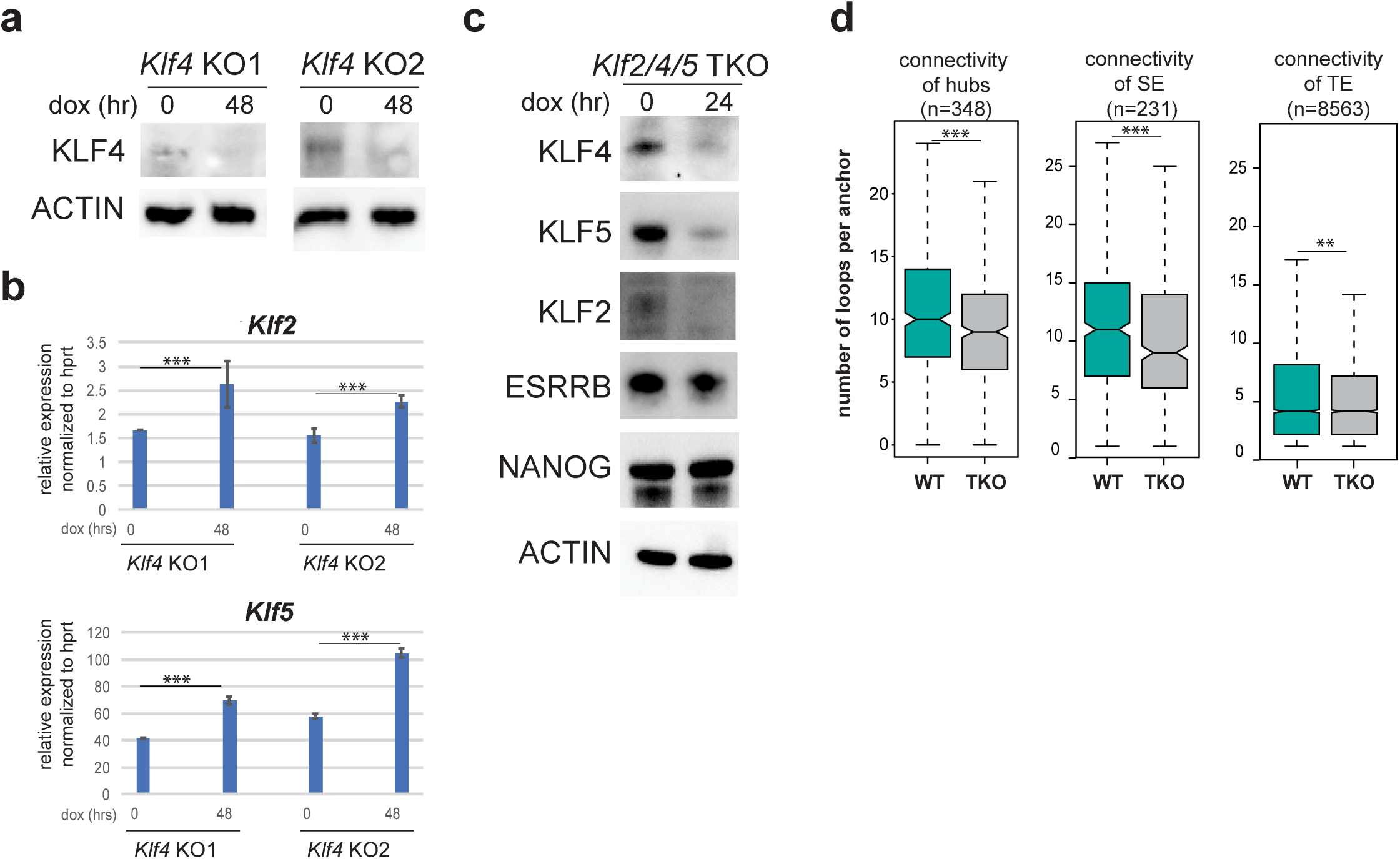
**a**, Western blot analysis showing KLF4 protein levels before (0) and after (48hr) dox-induction in two ESC clones that harbor dox-inducible CRISPR-Cas9 and gRNAs that target the *Klf4* gene (KLF4 KO1 and KLF4 KO2). **b**, RT-qPCR showing elevated levels of *Klf2* and *Klf5* genes in dox-induced KLF4 KO ESCs. **c**, Western blot showing levels of indicated proteins in a clonal population of ESCs containing an inducible CRISPR-Cas9 construct and gRNAs that target the *Klf2*, *Klf5* and *Klf4* genes. Cells were either untreated (0, wild type or WT cells) or treated with dox for 24 hours (triple knock-out or TKO). **d**, Boxplot showing the connectivity of H3K27ac HiChIP anchors that contain hubs, supoerenhancers (SE) or typical enhancers (TE) in WT or TKO ESCs. Asterisks indicate significance as calculated using Wilcoxon rank sum test.

**Supplementary Figure 7.**
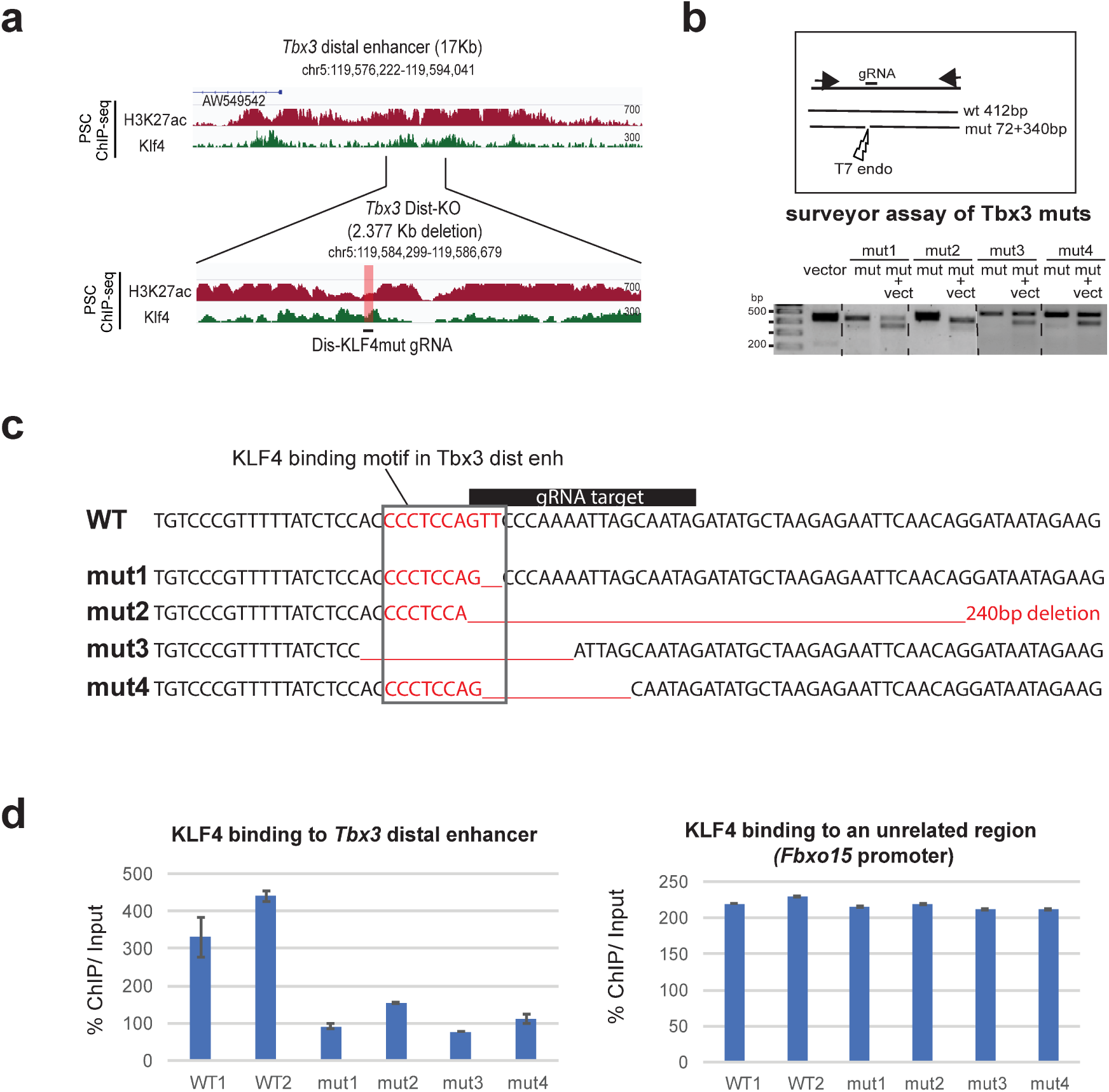
**a**, IGV tracks of H3K27ac and KLF4 ChIP-seq in PSCs showing the whole *Tbx3* distal enhancer (top), the region that was deleted by CRISPR/Cas9 (Dist-KO, bottom, see Fig.4f) and the location of the gRNA used to mutate a specific KLF4 binding motif (Dis-KLF4mut gRNA). **b**, Genotyping strategy of the surveyor assay used to detect mutation/indel at the target KLF4 binding site within the distal *Tbx3* enhancer (Dis-KLF4mut). The results for 4 homozygously mutated clones (mut1-4) are shown. **c**, Sequencing results of the four Mut clones compared to the wild type (WT). **d**, ChIP-qPCR showing the relative levels of KLF4 binding to *Tbx3* distal enhancer in two WT clones and four Mut clones (left panel). Values show percentage of ChIP signal over input. As control, binding of KLF4 to an unaffected region (*Fbxo15* promoter) was tested (right panel).

